# Repurposing Romidepsin for Osteosarcoma: Screening FDA-Approved Oncology Drugs with Three-Dimensional Osteosarcoma Spheroids

**DOI:** 10.1101/2025.05.22.654354

**Authors:** Emily E. Seiden, Spencer M. Richardson, Leah A. Everitt, Gabrielle J. Knafler, Gavin P. Kinsella, Alyssa L. Walker, Venetia A. Whiteside, James D. Buschbach, Deep A. Gandhi, M. Reza Saadatzadeh, L. Daniel Wurtz, Patrick J. Getty, Sheldon L. Padgett, Rance M. Gamblin, Michael O. Childress, Christopher M. Fulkerson, Karen E. Pollok, Christopher D. Collier, Edward M. Greenfield

**Author notes:** Corresponding Author: Edward M. Greenfield PhD; Department of Orthopaedic Surgery, Indiana University School of Medicine, MS 371 ORTS, Indianapolis, IN 46202, USA. these authors contributed equally to this work (Christopher D. Collier, Edward M. Greenfield). One or more of the authors has received funding from the Merilyn Hester Scholarship (EES), Leon Kenneth Knoebel Fellowship (EES), NIH T32CA27370 (EES), the Orthopaedic Research Education Fund Resident Clinician Scientist Training Grant OREF-RCSTG (CDC), the Dudley P. Allen Fellowship (CDC), NIH R21CA209304 (EMG), Indiana Univ Simon Cancer Center EDT/TMM Pilot Award (EMG, KEP, CDC), Big Ten Cancer Research Consortium Foundation Kenneth and Verna Mae Jessen Award (EMG, KEP, MOC, CMF), Strides For Sarcoma (EMG), Research Grant from the Rally Foundation for Childhood Cancer Research (EMG), Angie Fowler AYA Cancer Research Fund Pilot Program Grant (EMG), DOD Impact Award: CA210123 (KEP, MRS), DOD Impact Award: CA230160 (KEP, MRS), P30 CA082709 (KEP, MRS), The Danaher Foundation (KEP, MRS), The Caroline Symmes Children’s Cancer Endowment (KEP, MRS), and The Tyler Trent Cancer Research Endowment (KEP, MRS).

## Abstract

Osteosarcoma is the most common primary malignant bone tumor and predominantly affects children, adolescents, and young adults. It is the third most common cause of cancer-related deaths among 9-24-year-olds. Despite aggressive chemotherapeutic and surgical therapies, the survival rate is only 25% for patients with detectable lung metastases at diagnosis and only 70% in patients that present without detectable lung metastases. The poor prognosis is due to growth of metastases irrespective of whether they are initially large enough to detect clinically. It is therefore necessary to develop new methods to target the growth of lung micrometastases. An NCI panel of FDA-approved oncology drugs was therefore screened using three highly metastatic human osteosarcoma cell lines. To more closely approximate *in vivo* micro-metastases, the screen used a 3D multicellular *in vitro* osteosarcoma spheroid (sarcosphere) model. Among 13 hits from the initial screen, we identified the histone deacetylase inhibitor (HDI) romidepsin as the most promising inhibitor in secondary screens based on sarcosphere viability. Romidepsin potency was evident with and without standard-of-care chemotherapeutics (MAP: Methotrexate, Adriamycin, Cisplatin) at drug concentrations that are clinically achievable and did not affect non-transformed cells. By those criteria, romidepsin also substantially outperformed the other three FDA-approved HDIs and eight HDIs in clinical trials. Importantly, sarcospheres derived from 30-50% of human and canine patient samples were also sensitive to romidepsin with ED50s 10- to 700-fold less than the Cmax in human patients. Based on these 3-D screening approaches, romidepsin is a promising drug to repurpose for osteosarcoma.

**Significance Statement:** Our unbiased sarcosphere-based drug screen identified romidepsin as a promising candidate to repurpose for canine and human patients with metastatic osteosarcoma. This screening strategy allowed us to identify romidepsin-sensitive and -resistant patients. Sarcosphere-based screening may therefore be useful to stratify patients most likely to respond clinically to romidepsin or other drugs.

## Introduction

Osteosarcoma is the most common primary malignant bone tumor and is the 3^rd^ most common cause of death in pediatric cancer patients ages 9-24^1, 2^. With surgery and standard-of-care chemotherapy (consisting of MAP: high-dose methotrexate, doxorubicin, cisplatin), the five-year survival rate of osteosarcoma patients is only 70%, which has not improved since the 1980s^2^. For patients with detectable lung metastases, the survival rate is only 25%^1, 3^. Lethality in osteosarcoma is due to growth of lung metastases irrespective of whether they are detectable at diagnosis^3, 4^. Novel therapeutic agents targeting the growth of both clinically detectable lung metastases and undetectable lung micrometastases are therefore needed to improve outcomes in osteosarcoma. Developing new therapeutics to treat osteosarcoma is challenging for many reasons. Osteosarcoma is genetically complex with substantial heterogeneity between individual patients^5–8^. It is therefore unlikely that a single agent will be successful for all patients and personalized treatments may be necessary. Additionally, osteosarcoma is a rare disease, with about 800 new diagnoses per year in the United States, which makes clinical trials difficult and severely reduces commercial interest^9^. Repurposing existing FDA-approved drugs is therefore a promising alternative as it reduces the time and cost of drug discovery and provides an opportunity to quickly advance therapies with known safety profiles into clinical trials^10^. This study therefore screened the NCI (National Cancer Institute) panel of FDA-approved oncology drugs to identify promising agents to repurpose for osteosarcoma. 3D cell culture models more closely represent the *in vivo* tumor environment compared with traditional 2D monolayer cultures, and better correlate with *in vivo* responses to chemotherapeutics^11–15^. Advantages of 3D models include cell to cell interactions and biological gradients of oxygen, nutrients, and metabolic waste products^11, 16–22^. 3D models also have genetic profiles and growth kinetics that are more similar to *in vivo* microtumors^11–15^. This study therefore used a highly-uniform 3D osteosarcoma spheroid (sarcosphere) platform that we previously developed^23^ to screen the National Cancer Institute (NCI) panel of FDA-approved oncology drugs. Importantly, the platform has acceptable throughput as we routinely obtain hundreds of sarcospheres from a single preparation of cells. Screening was performed with and without MAP chemotherapy with sarcospheres derived from three different highly metastatic human osteosarcoma cell lines. The purpose of this unbiased screen was to identify agents that were potent against sarcospheres at clinically achievable doses, additive-to-synergistic with MAP chemotherapy, and non-toxic on normal human osteoblasts and non-transformed small airway epithelial cells. Lung cells were included because intranasal therapy is being developed to administer therapeutics directly to lung metastases of osteosarcoma patients^4, 24, 25^. Romidepsin, the most promising drug based on those criteria, was also tested on sarcospheres obtained from low-passage patient-derived cells. Both human and canine patients were included because osteosarcoma is biologically similar in both species^24, 26, 27^. For example, canine and human osteosarcomas have similar histology and disease progression and show similar responses to standard therapies^28^. In addition, gene expression between human and canine cell lines and pre-treated primary tumor samples is similar, indicating that canine osteosarcoma can be used as a model for disease progression^29^. In addition, translational studies are facilitated as osteosarcoma is more common in canine patients and canine osteosarcoma patients have a survival rate of less than 10%^30^. The overall layout of our study, including drug screen and hit discovery, hit confirmation and characterization, and evaluation of the most promising drug is summarized in Figure 1.

**Figure 1.**
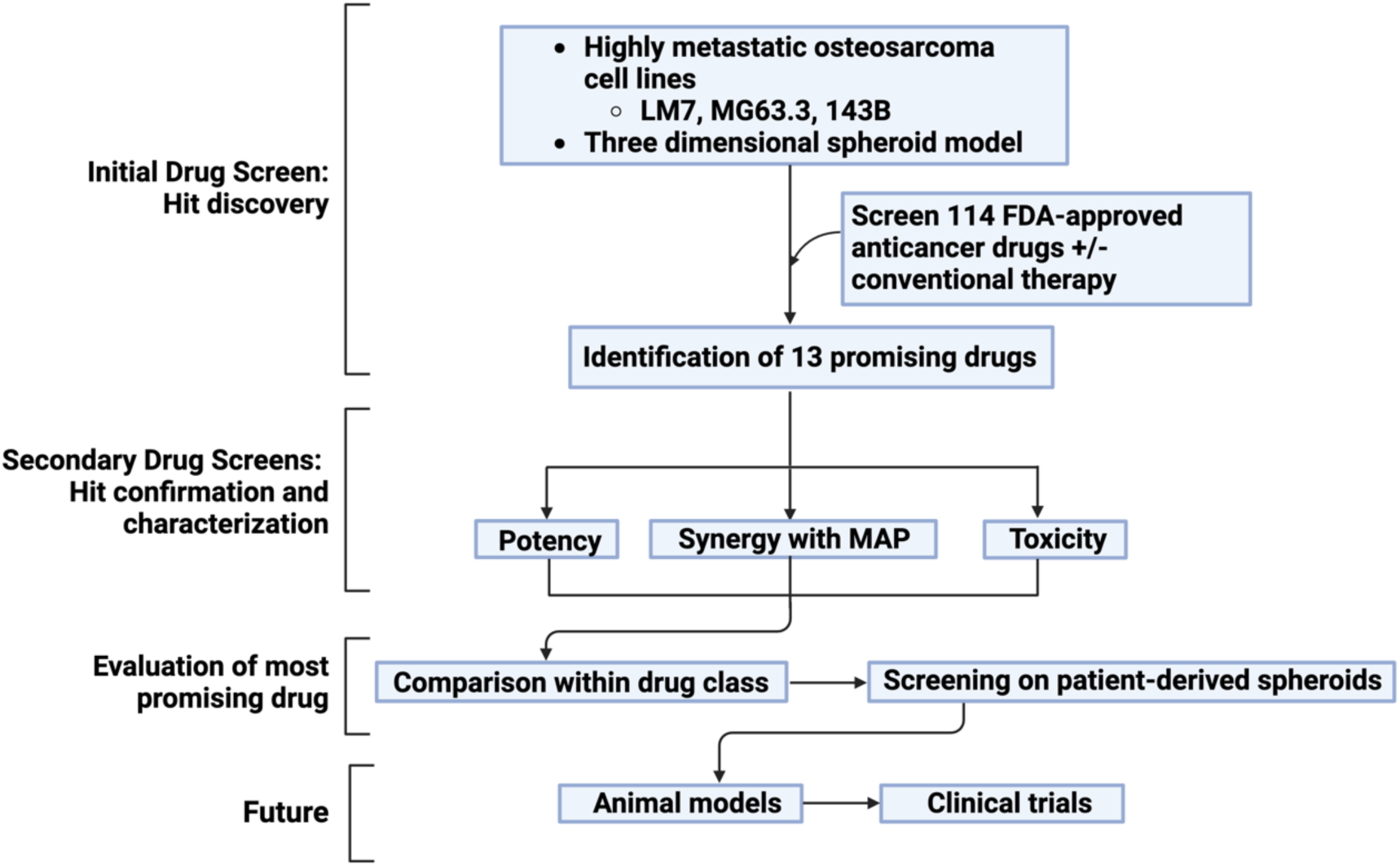
Repurposing promising therapeutics study design. Drug screen used three highly metastatic osteosarcoma cell lines in a 3D sarcosphere platform. The NCI panel of FDA-approved oncology drugs was screened with and without conventional MAP therapy and 13 drugs were identified. The 13 hits were evaluated for potency on sarcospheres with and without MAP therapy and for toxicity on NHOst and SAEC cells. The most promising drug, romidepsin, was compared to a panel of HDIs that were not included in the initial screen. To further evaluate the top hit, romidepsin was screened on a variety of human and canine patient-derived sarcospheres. Future directions include animal models and canine and human clinical trials. Figure 1 was created with Biorender.com.

## Materials and Methods

### Cell Culture

Three well characterized highly metastatic osteosarcoma cell lines were used in this study^31^. The 143B cell line (RRID: CVCL_2270)^32^, derived from the HOS-TE85 cell line (RRID:CVCL_0439) was obtained from the American Type Culture Collection (Manassas, VA, USA). The MG63.3-GFP cell line (RRID:CVCL_WL01)^33^, derived from the MG63 cell line (RRID:CVCL_0426) was obtained from the laboratory of C. Khanna DVM, PhD (National Cancer Institute, Bethesda, MD, USA). The LM7 cell line (RRID:CVCL_0515)^34^, derived from the SaOS-2 cell line (RRID:CVCL_0548), was obtained from the laboratory of E.S. Kleinerman, MD (MD Anderson Cancer Center, Houston, TX, USA). Small Airway Epithelial Cells (SAEC, ATCC, Catalog #PCS-301-010) and Normal Human Osteoblast Cells (NHOst, Lonsa, Catalog #CC-2538) were used for toxicity screening. All experiments were performed on cells passaged 3-7 times after thawing from liquid nitrogen. Cell culture media was advanced minimum essential medium (Gibco, Catalog #12492013,) supplemented with 10% fetal bovine serum (Cytiva Catalog #SH30071.03), 1% Glutamax (Gibco, Catalog #35050-06), and either 1% penicillin-streptomycin (Gibco, Catalog #15140122) or 1% antibiotic-antimycotic (Corning Catalog #30-0004-CL). Medium was sterile filtered before use. All cell cultures were maintained at 37°C in a humidified 5% CO_2_ incubator.

### Isolation of Primary Cells from Patient Tumors

Osteosarcoma tissue was obtained during standard-of-care procedures (biopsies or resections) in human patients treated at Indiana University Health and Riley Children’s Hospital with Institutional Review Board (IRB) approval (IRB #1501467439). Tissue from canine osteosarcoma patients was obtained from standard-of-care amputations at the Purdue University College of Veterinary Medicine and the Metropolitan Veterinary Hospital (Akron, Ohio). Osteosarcoma diagnosis was confirmed by review of pathology reports. Primary osteosarcoma cells were isolated by collagenase/mechanical dispersion using a modification of a previously described protocol^35^. Demographic and clinical information for human and canine patients are in Tables 1 and 2. Briefly, tissue samples were collected in CO_2_ independent media (Gibco, Catalog #18045-088) supplemented with 1% GlutaMax (Gibco Catalog #35050-06) and 1% antibiotic-antimycotic (Corning, Catalog #30-0004-CL) and placed on ice during transportation. The tissue was rinsed with cold PBS (Cytiva, Catalog #SH30028.02) and minced prior to incubation in 10 mL of culture media containing collagenase Type II (750 units/mL with 13-18 units of clostripain/mL, Worthington, Catalog #LS004176) for 3 hours at 37°C and 5% CO_2_. The media and tissue fragments were centrifuged at 200g for 5 minutes. Pellets with a visible layer of red blood cells were then treated with 5 mL of ACK Lysing Buffer (Gibco, Catalog #A10492-01) for 1 minute and then centrifuged at 200g for 2 minutes. Pellets were resuspended in culture media, pipetted up and down 15 times to disperse tissue fragments, and centrifuged at 200g for 5 minutes. Finally, any remaining tissue fragments were crushed with a 25 mL pipette and transferred with released cells and culture media to a 100 mm tissue culture dish and incubated at 37°C and 5% CO_2_. Media was changed every three days and the cells were passaged to 150 mm dishes at 5000 cells/cm^2^ when a confluence of ∼80% was reached. Aliquots of the single-passage patient-derived osteosarcoma cells were frozen in 10% DMSO (Sigma-Aldrich D2650), 40% FBS (Cytiva SH30071.03), and 50% culture media and stored in liquid nitrogen. The TT2-77 xenoline is a patient cell line that was isolated from a patient-derived xenograft (PDX) that was previously developed and characterized by the Pollok lab^5^.

**Table 1.**
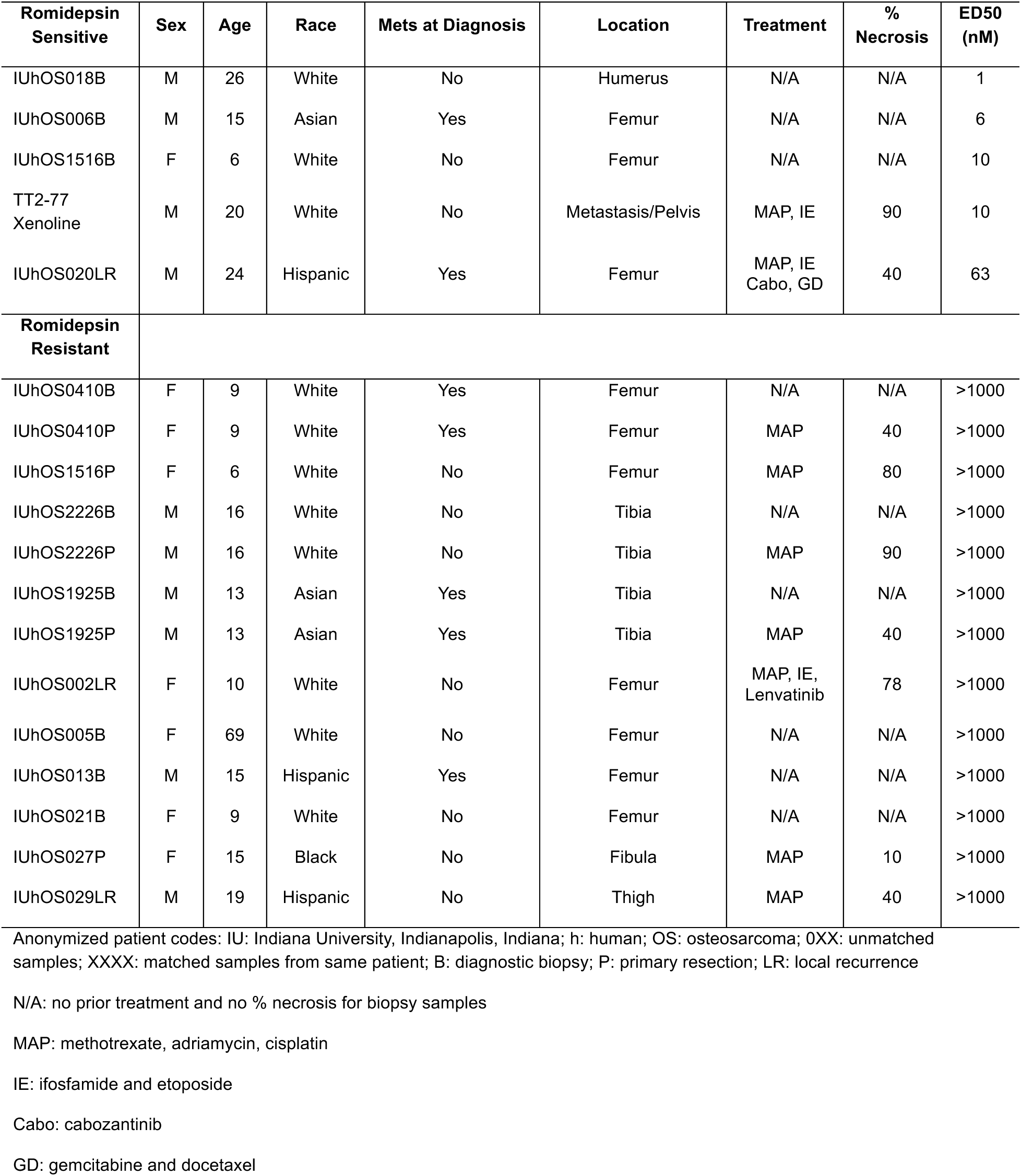
Clinical Information for Human Patients and Romidepsin ED50s.

**Table 2.** Clinical Information for Canine Patients and Romidepsin ED50s.

| Romidepsin Sensitive | Sex | Age | Breed/Breed Classification | Tumor Location | ED50 (nM) |
| --- | --- | --- | --- | --- | --- |
| MVHcOS001 | M | 5y2mo | Bernese Mountain Dog/Giant | Right distal radius | 6 |
| MVHcOS002 | M | 6y2mo | Great Pyrenees/Giant | Right distal tibia | 22 |
| PUcOS003 | F | 6y8mo | Greyhound/Large | Right tibia | 46 |
| PUcOS004 | M | 9y4mo | Greyhound/Large | Left humerus | 57 |
| PUcOS005 | M | 8y8mo | German Shepherd/Large | Distal radius | 65 |
| **Romidepsin Resistant** |  |  |  |  |  |
| MVHcOS006 | F | 5y9mo | Cane Corso/Giant | Left proximal humerus | >1000 |
| MVHcOS007 | F | 4y6mo | Mastiff Mix/Large | Right distal radius | >1000 |
| MVHcOS008 | F | 7y11mo | German Shepherd/Large | Right distal radius | >1000 |
| PUcOS009 | M | 11y4mo | Mixed Breed/Unknown | Right proximal femur | >1000 |
| MVHcOS010 | F | 10y4mo | Alaskan Malamute/Large | Right distal radius | >1000 |

Anonymized patient codes: MVH: Metropolitan Veterinary Hospital, Akron, Ohio; PU: Purdue University, West Lafayette, Indiana; c: canine; OS: osteosarcoma; XXX: sample number (all from amputations)

### 3D *in vitro* Sarcosphere Generation

Sarcospheres were generated (1 sarcosphere/well in 96 well plate) using the technique we developed previously^23^ that was modified from a technique described for epithelial cancers^19^. Sarcospheres of the desired size to generate similar oxygen and nutrient gradients to an *in vivo* tumor (400 um in diameter)^19^ were obtained by adjusting the number of cells per well as we previously described (143B: 2500 cells/sarcosphere, MG63.3-GFP: 1000 cells/sarcosphere, LM7: 8000 cells/sarcosphere)^23^. The number of patient-derived cells required to obtain sarcospheres with 400 um diameters was determined for each patient and varied from 1000-15,000 cells per sarcosphere. For that purpose, sarcosphere sizes were measured using the Incucyte S3 Live Cell Analysis System (Sartorius, Gottingen, Germany). Brightfield images were captured using a 4x or 10x objective. Incucyte analysis software was used to automatically detect area of sarcospheres in 96 well plates and diameters were calculated using the formula for area of a circle.

Round bottom 96-well plates were coated with 50 µL/well of 0.5% poly 2-hydroxyethyl methacrylate (poly-HEMA) (Polysciences Inc., Catalog #09689-25) in 100% ethanol, and incubated at 37°C for 3 days before use to obtain a nonadherent surface. Alternatively, pre-coated Nunclon Sphera-Treated plates (Nunc, Catalog #174929) or SBio PrimeSurface Ultra-low Attachment plates (SBio, Catalog #MS-9096UZ) were used. Highly metastatic osteosarcoma cells cultured in monolayers at 50-80% confluence were lifted with Accutase (Innovative Cell Technologies, Inc., Catalog #NC9839010), resuspended in media and diluted to the desired concentration. Cell suspensions were combined with 2.5% Matrigel (Corning, Catalog #354234) and 100 µL was added to each well. Three to six wells of the 96-well plate contained 100 µL of 2.5% Matrigel and media without cells to provide background values. The peripheral wells of each plate contained PBS or sterile water to minimize evaporation in wells with sarcospheres. Following cell seeding, plates were centrifuged (1000 g for 10 minutes at 4°C) and then were incubated at 37°C, 5% CO_2_. Sarcospheres were matured for 24 hours prior to drug treatment (Day 0). Then, 100 µL drug treatment or vehicle was added to each well. Plates were incubated for 48 hours and analyzed (Day 2). Resazurin reduction on Day 2 by vehicle-treated sarcospheres were compared to resazurin reduction on Day 0 to calculate sarcosphere growth without drug over 2 days in selected experiments.

### Chemotherapeutics

The panel of 114 FDA-approved oncology drugs (NCI Panel #4804082 and #4803082) was obtained from the National Cancer Institute (NCI) Developmental Therapeutic Program at 10 mM in DMSO and stored at -80°C until use. Storage information for the 13 drugs used in the secondary screen is listed in Supplemental Table 1. Romidepsin and all other HDIs (Selleck Chemicals) were dissolved in DMSO and stored at -80°C until use. Cisplatin (Tocris Biosciences, Bristol, UK) was solubilized in saline to avoid inactivation by DMSO^36^. Doxorubicin and methotrexate (Tocris Biosciences, Bristol, UK) were solubilized in water or PBS respectively, as done in previous studies from this laboratory^37^. Doses of all drugs were obtained following serial dilution from stock solutions. All wells in an experiment received the same concentration of the relevant vehicle.

### Resazurin Reduction Assay

Viability of sarcospheres was assessed by resazurin reduction measurements of metabolic activity. Resazurin (Sigma-Aldrich, Catalog #R7017-1G) was dissolved in PBS at 0.15 mg/mL. 20 µL of resazurin solution was added to each well containing sarcospheres in 200 µL of media and to the background wells without sarcospheres. Plates were incubated at 37°C in 5% CO_2_ for 2 or 6 hours. Fluorescence readings were taken at an excitation wavelength of 535nm and an emission wavelength of 590nm (Tecan Genios Pro, Mannedorf, Switzerland). The average fluorescence from the background wells was subtracted from vehicle and treatment wells.

### Initial Screen of FDA-approved Oncology Drugs

Sarcospheres were matured for 24 hours, then treated for 48 hours with media containing 10 µM of each drug and 0.1% DMSO. This relatively high drug concentration was selected to minimize false negatives. Control and background wells also contained 0.1% DMSO. Each drug was tested in duplicate non-adjacent wells to minimize the impact of well position and edge effect. All drugs were tested with and without MAP on sarcospheres derived from the three highly metastatic cell lines LM7, 143B, and MG63.3-GFP. Effects on resazurin reduction are reported as growth inhibition as previously described by the NCI^38^ and used by our group^23^. Growth inhibition represents a more clinically translatable measure for screening studies wherein a value of 0% represents no tumor growth since start of treatment, 0-100% represent tumor growth where 100% is the untreated control, and values less than 0% represent tumor reduction where -100% represents no viable tumor cells. Drugs from the NCI panel were screened with and without MAP chemotherapeutics. Methotrexate, doxorubicin, and cisplatin were dosed to inhibit sarcosphere growth by 20% by combining the individual chemotherapeutics at a fixed ratio of their individual GIC50s (growth inhibition of 50%). This lower concentration of MAP was selected to maintain sufficient dynamic range to identify hits in the screen. Concentrations of MAP for the initial screen are listed in Supplemental Table 2. Screening assay quality was assessed throughout by direct examination of duplicate data which remained highly consistent for all drugs and conditions tested and by calculating Z’ factors, a measure of assay quality in high-throughput screening used to assess assay signal dynamic range and variability^39^. The Z’ factor threshold for an “excellent” assay (0.5 to 1.0) was met for all individual plates and the mean Z’ factor for all plates combined was 0.70 with a standard deviation of 0.01.

### Secondary Screens of Promising Drugs

FDA-approved drugs in the top quartile by efficacy of our initial screen, as defined by mean growth inhibition across all cell lines with and without MAP were considered for inclusion in the secondary screens. An exhaustive literature review was performed for each drug considered. Priority was given to drugs that performed best in the screen across all conditions at clinically achievable concentrations, had unique mechanisms of action, demonstrated efficacy in other solid cancers, had favorable or untested efficacy in osteosarcoma basic research or clinical trials, and were not already recommended as first- or second-line therapies for patients with osteosarcoma by the National Comprehensive Cancer Network. This process identified 13 promising drugs for the secondary screens evaluating potency, toxicity, and potential for combination therapy with MAP. For these secondary analyses, we reported drug responses as percent of control wherein 0% represents a complete response and 100% represents no response similar to the untreated control. This analysis allowed us to more accurately pool results from 3-6 independent experiments as values do not fluctuate based on viability at the start of treatment. Potency experiments were performed on sarcospheres following serial dilution of each drug at log-based concentrations from 0.001 µM to 10 µM to determine the ED_50_ (effective dose at which viability is 50% of the control).

Combination therapy experiments were performed by comparing drug response across multiple doses with and without MAP at a fixed dose. MAP was dosed to obtain 50% reduction in resazurin reduction by combining each individual chemotherapeutic at a ratio of their individual ED50s. Concentrations of MAP used in secondary screen are listed in Supplemental Table 2. Toxicity experiments were performed on monolayers of non-transformed NHOst and SAEC cells at 50-80% confluency in similar fashion to calculate the TD_50_ (toxic dose at which viability is 50% of the control).

### Statistical Analysis

For the initial screen of FDA-approved drugs, unsupervised hierarchical clustering of cell lines based on drug responses were used to generate a heatmap and a sample dendrogram. Based on the mean response the drugs were sorted. R-project (version 3.1.2.) hclust function with Ward-D2 method was used to generate the dendrogram. Dendrogram at bottom illustrates the arrangement of the cell lines based on comparison of responses via hierarchical clustering. The height of the tree branch connecting end nodes (cell lines) in the dendrogram is proportional to the value of the intergroup dissimilarity. The color bar on the left of the heatmap shows the classification of the drugs.

All other data are presented as a mean of at least 3 independent experiments. Error bars represent standard deviations. Each experiment included a minimum of 3 sarcospheres per group. ED50s and TD50s were calculated using a 3-parameter logistic regression analysis (GraphPad Prism version 10.2.3). One-way ANOVA with Dunnett’s multiple comparisons test (GraphPad Prism version 10.2.3) was used to compare treated groups to control in dose response experiments. Two-way ANOVA with Sidak’s multiple comparisons test (GraphPad Prism version 10.2.3) was used to compare treated groups to control groups in preliminary screen of other HDIs (FDA-approved and in clinical trials). Heat maps were ranked by the means of each row in the heat map (highest to lowest value). Blue indicates the highest value of a heat map, red is the lowest, and white indicates the median of the row means. Synergy scores for Bliss and Loewe models of additivity were calculated using SynergyFinder+^40^. All graphs and heat maps were generated using GraphPad Prism (version 10.2.3).

### Data Availability Statement

Data were generated by the authors and available upon request from the corresponding author.

## Results

### HDI, proteosome inhibitors, and topoisomerase inhibitors were most effective in initial screen

In the initial unbiased screen, the NCI-panel of FDA-approved oncology drugs was tested at 10 μM against sarcospheres derived from three highly metastatic human osteosarcoma cell lines (143B, MG63.3-GFP and LM7). Dark blue in heatmap indicates inhibition of sarcosphere growth based on resazurin reduction (Figure 2A). Unsupervised clustering analysis showed that 143B and MG63.3-GFP sarcospheres responded relatively similarly, while responses by LM7 sarcospheres were relatively distinct (Figure 2A). Histone deacetylase inhibitors (HDI), proteosome inhibitors, and topoisomerase inhibitors most effectively reduced sarcosphere growth both with and without MAP chemotherapeutics in sarcospheres generated from all three osteosarcoma cell lines (Figures 2A-B). Few drugs from the other drug classes reduced viability (Figure 2B). In agreement with our previous study of the osteosarcoma standard-of-care chemotherapeutics^23^, doxorubicin was effective with sarcospheres derived from all three cell lines and methotrexate was effective with 143B sarcospheres. In contrast, cisplatin had no detectable effect on sarcospheres derived from any of the cell lines, presumably because it was inactivated during preparation of the NCI panel by solubilization in DMSO^36^.

**Figure 2.**
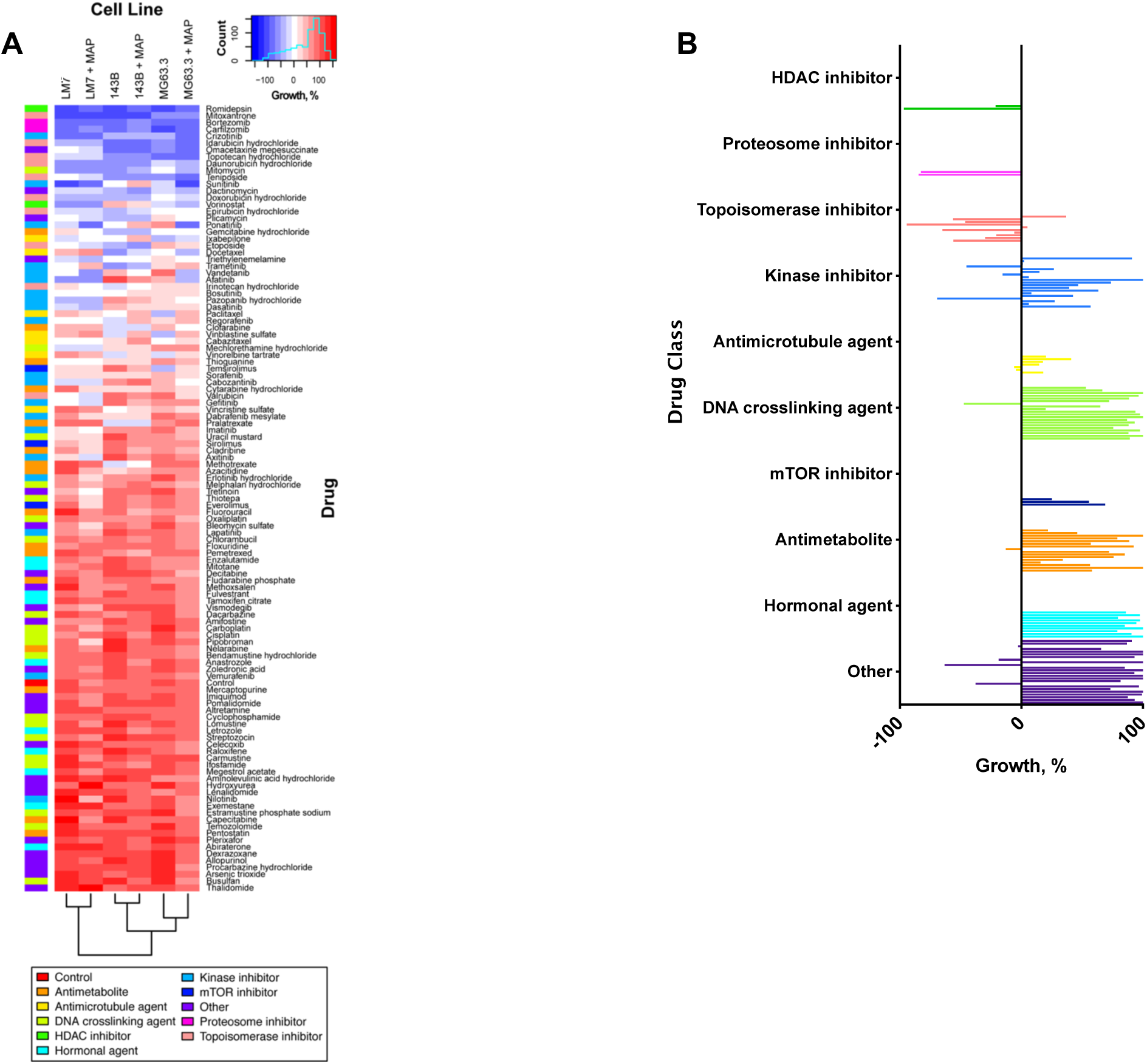
Top hits of initial screen are histone deacetylase inhibitors, proteosome inhibitors, and topoisomerase inhibitors. **(A)** Screen of FDA-approved oncology drugs on sarcospheres generated from 143B, MG63.3-GFP, and LM7 cell lines. Heat map shows inhibition of growth in blue and little or no effect in red. Colored bars on left of figure indicate control or drug class. Results of unsupervised clustering analysis are shown at the bottom of the figure. Sarcospheres were screened with the individual drugs (10 *μ*M) both with and without MAP chemotherapeutics for 48 hours. All wells contained 0.1% DMSO as vehicle. Growth was assessed by change in resazurin reduction during the 48 hours without drug. A growth percentage value of 0 represents no tumor growth since the start of treatment; values 0-100 represent tumor growth where 100 is the untreated control; values less than 0 represent tumor reduction where -100 represents no viable tumor cells. **(B)** Summary of the initial screen showing the mean of the rows for each drug in panel A.

### Secondary screens identified the HDI romidepsin as the most promising drug to repurpose for osteosarcoma

After exclusion of the standard-of-care doxorubicin and its analogues (idarubicin, daunorubicin, and epirubicin), 13 of the remaining 22 most effective drugs in the initial screen were selected for secondary screening. Priority was given to drugs that performed best in the screen across all conditions at clinically achievable concentrations, had unique mechanisms of action, demonstrated efficacy in other solid cancers, had favorable or untested efficacy in osteosarcoma basic research or clinical trials, and were not already recommended as first- or second-line therapies for patients with osteosarcoma by the National Comprehensive Cancer Network. The 13 drugs included both HDIs (romidepsin and vorinostat) and both proteosome inhibitors (bortezomib and carfilzomib) in the initial screen as well as inhibitors of tyrosine kinases (afatinib, crizotinib, ponatinib, and vandetinib), topoisomerase (mitoxantrone and teniposide), transcription (plicamycin) and translation (omacetaxine), and a DNA crosslinker (mitomycin C). Dose-response experiments confirmed the initial screening results for all 13 drugs (Supplemental Figure S1A). Bortezomib had the lowest ED50 with 143B and MG63.3-GFP sarcospheres, while romidepsin had the lowest ED50 with LM7 sarcospheres and the second and third lowest with MG63.3-GFP and 143B sarcospheres, respectively (Supplemental Figure S1B-D and Supplemental Table 3). The ED50s were also compared to the clinically-achievable level (Cmax) of each drug, listed in Supplemental Table 3^41–45^, which represents the peak plasma concentration measured and tolerated from a single administration. Carfilzomib had the highest overall mean of the three Cmax/ED50 ratios and romidepsin ranked second (Figure 3A). A comparison of the Cmax/ED50 ratios of sarcospheres from each cell line showed that romidepsin had the highest ratio with LM7 sarcospheres, the second highest ratio with MG63.3-GFP sarcospheres, and the fifth highest with 143B sarcospheres (Figure 3A).

**Figure 3.**
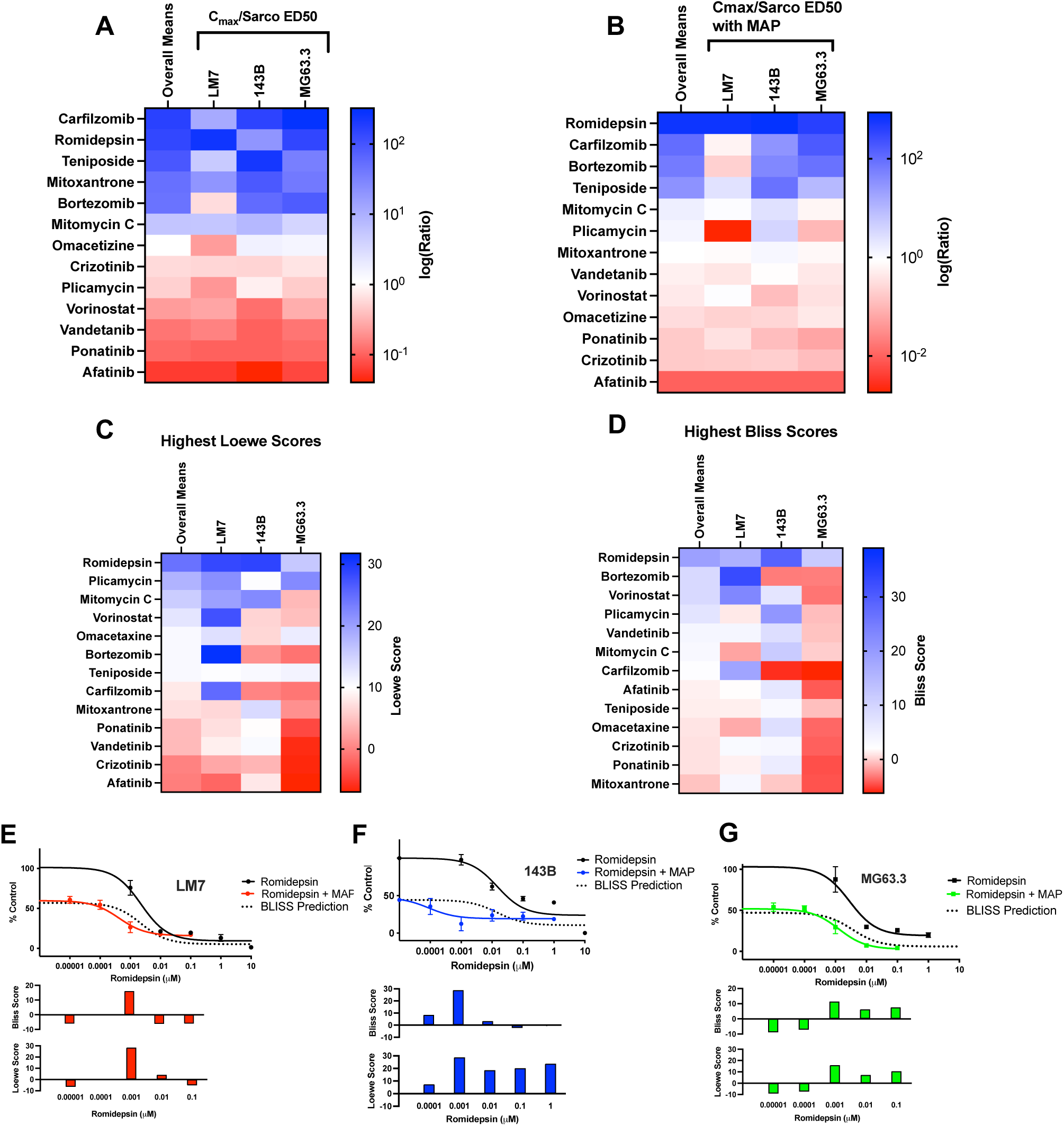
The HDAC inhibitor romidepsin is most promising of the 13 FDA-approved oncology drugs in the secondary screen. All heat maps in this Figure are sorted by the overall row means shown in the first column of each heat map. Dark blue indicates highest values (most favorable), dark red the lowest values (least favorable), and white is set at the median of the row means. **(A-B)** Ratios of Cmax/sarcosphere ED50 with and without MAP reflect potency at clinically-achievable concentrations. Cmax’s and ED50s are shown in Supplemental Table 3 and Supplemental Figure S1. ED50s were determined for sarcospheres derived from LM7, 143B, and MG63.3-GFP cell lines in N=3 independent dose response experiments with each drug. Each experiment included at least 3 sarcospheres per group. **(C-D)** Synergy scores in combination with MAP. Highest synergy scores at concentrations less than Cmax were calculated for each drug using the Bliss and Loewe models of additivity from the data shown in panels E-G and Supplemental Figures S2-4. **(E-G)** Romidepsin dose responses with and without MAP for sarcospheres derived from LM7, 143B, and MG63.3-GFP cell lines. Solid black lines indicate romidepsin dose responses without MAP, colored lines indicate romidepsin dose response with MAP. Dotted lines indicate Bliss prediction of perfect additivity. Bar graphs show synergy scores at each individual drug concentration.

The 13 drugs were also screened in combination with the three MAP chemotherapeutic drugs because combination therapy with MAP is a likely application for any new osteosarcoma drug. When ED50s were measured in the presence of MAP, romidepsin had the most favorable Cmax/ED50 ratio in sarcospheres derived from all three cell lines (Figure 3B). Moreover, by the Bliss and Loewe additivity models^46^, romidepsin had the highest overall mean of synergy scores of the 13 drugs at concentrations less than Cmax (Figure 3C and D). In addition, romidepsin had additive-to-synergistic activity in combination with MAP in sarcospheres derived from all three cell lines. Additive-to-synergistic activity is indicated in Figure 3E-G by downward shifts in the romidepsin dose responses in combination with MAP compared with the Bliss additivity model depicted by the dotted lines. Importantly, the highest synergy scores for romidepsin were at 1 nM, which is substantially less than its Cmax of ∼700 nM (Figure 3E-G bar graphs). In comparison, the only other examples of additive-to-synergistic activity in combination with MAP by Bliss curve fitting were ponatinib and vorinostat with LM7 sarcospheres and plicamycin with 143B sarcospheres (Supplemental Figures S2-4).

To evaluate safety of the 13 drugs, we measured effects on viability of monolayers of normal human osteoblasts (NHOst) and non-transformed human small airway epithelial cells (SAEC) (Figure 4). We elected to use monolayers for this purpose rather than 3-D spheroids because neither osteoblasts nor SAECs form 3-D aggregates *in vivo*. SAEC were included because aerosol therapy is being developed to directly treat lung metastases of osteosarcoma patients^25, 47, 48^. Six therapeutic indices were calculated for each drug as the TD50 for NHOst and SAEC divided separately by each of the three sarcosphere ED50s. Romidepsin had the largest overall mean of the therapeutic indices on NHOst and SAEC cells (Figure 4). Romidepsin had 11-, 42-, and 162-fold higher NHOst therapeutic indices than the next highest drug in 143B, MG63.3-GFP, and LM7 sarcospheres respectively (Figure 4, Supplemental Figure S5A-D). Romidepsin also had the largest individual SAEC therapeutic index with LM7 sarcospheres and the second largest with 143B and MG63.3-GFP sarcospheres (Figure 4, Supplemental Figure S6A-D). Taken together, the potency and toxicity screens identified the HDI romidepsin as the most promising drug to repurpose for osteosarcoma. The next most promising FDA-approved drugs were bortezomib and carfilzomib, the only two proteosome inhibitors in the NCI panel.

**Figure 4.**
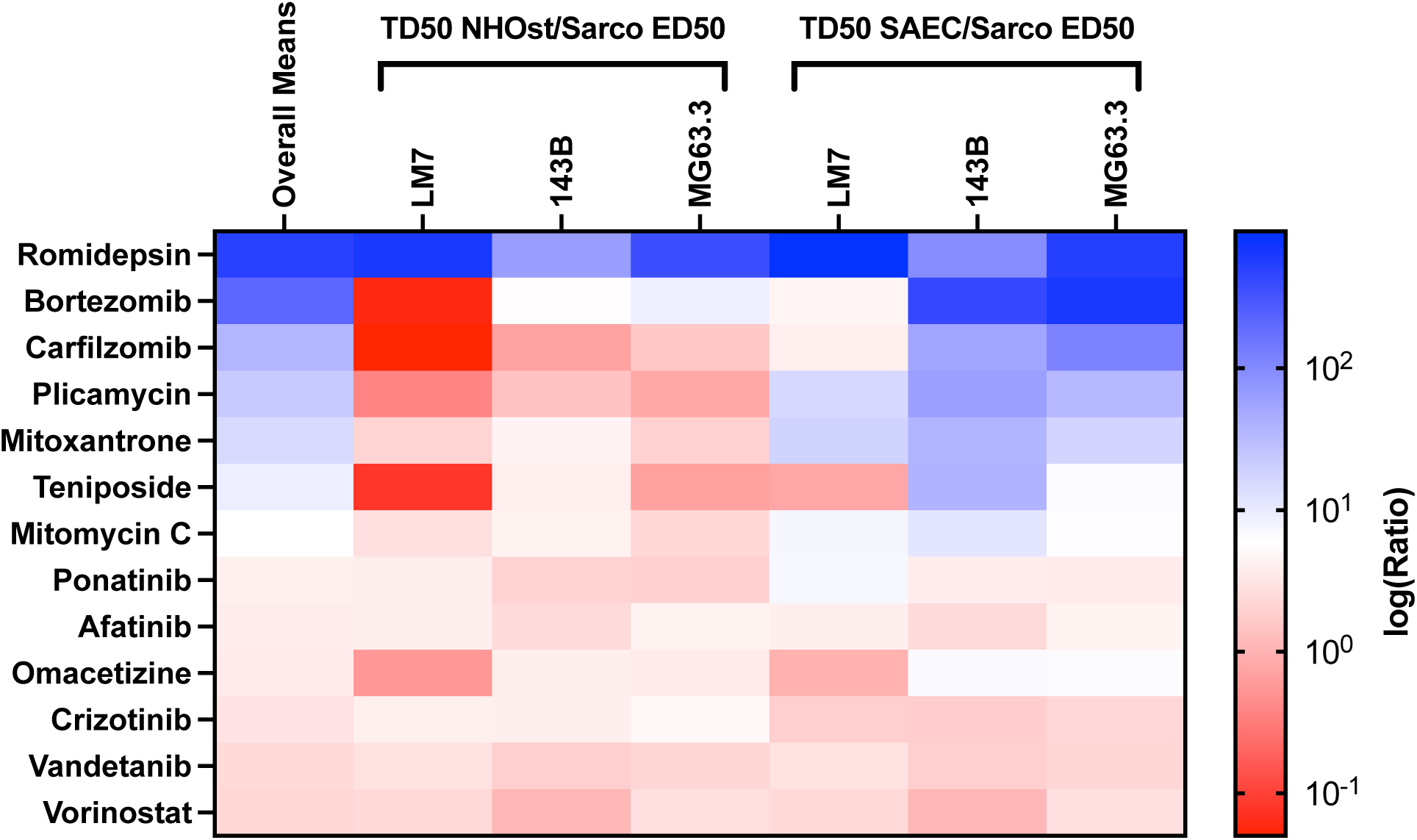
Romidepsin has the largest overall therapeutic index of all 13 hits from the initial screen. Data represents dose response experiments for top 13 hits from initial screen. Ratios of TD50/sarcosphere ED50 reflect therapeutic indices with normal human osteoblasts (NHOst) and non-transformed human small airway epithelial cells (SAEC). White on the color scale is set at the median of the overall row means, shown in the first column. TD50s, doses that reduce viability of NHOst and SAEC by 50%, are shown in Supplemental Figures S5A and S6A. The TD50/ED50 ratios are shown in Supplemental Figures S5B-D and S6B-D. TD50s were determined in N=3 independent dose response experiments with each drug. Each experiment included at least 3 culture wells per group. ED50s, doses that reduce viability of 143B, LM7, or MG63.3-GFP sarcospheres by 50% are shown in Supplemental Figure S1B-D and Supplemental Table 3.

### Romidepsin is more promising than all other tested HDIs

Each HDI has unique pharmacokinetics and pharmacodynamics, targets different classes of HDACs, and therefore could be more effective or safer than romidepsin^49–55^. Since only two HDIs were included in the initial screen, romidepsin was compared with the 3 other HDIs that are FDA-approved for oncology patients and seven that are in clinical trials. The HDIs in clinical trials were initially screened at their Cmax, with and without MAP chemotherapeutics, against sarcospheres derived from each of the osteosarcoma cell lines. AR-42 and resminostat were most effective in all of those preliminary studies (Supplemental Figure S7) and were therefore advanced for further study with the FDA-approved HDIs (belinostat, panobinostat, romidepsin, and vorinostat)). Dose response experiments with sarcospheres derived from all three cell lines showed that romidepsin had the lowest ED50s with and without MAP chemotherapeutics (Supplemental Table 3 and Supplemental Figure S8), the most favorable Cmax/ED50 ratios with and without MAP (Figure 5A-B), and the largest NHOst therapeutic indices (Figure 5C). Moreover, the effects of romidepsin and vorinostat in this screen are very similar to, and therefore confirm, the effects of those HDIs in the secondary screen (Supplemental Table 3). Therapeutic indices for the HDIs were not determined with the SAEC cells because they were less discriminatory than the NHOst therapeutic indices for the original 13 drugs (Figure 4).

**Figure 5.**
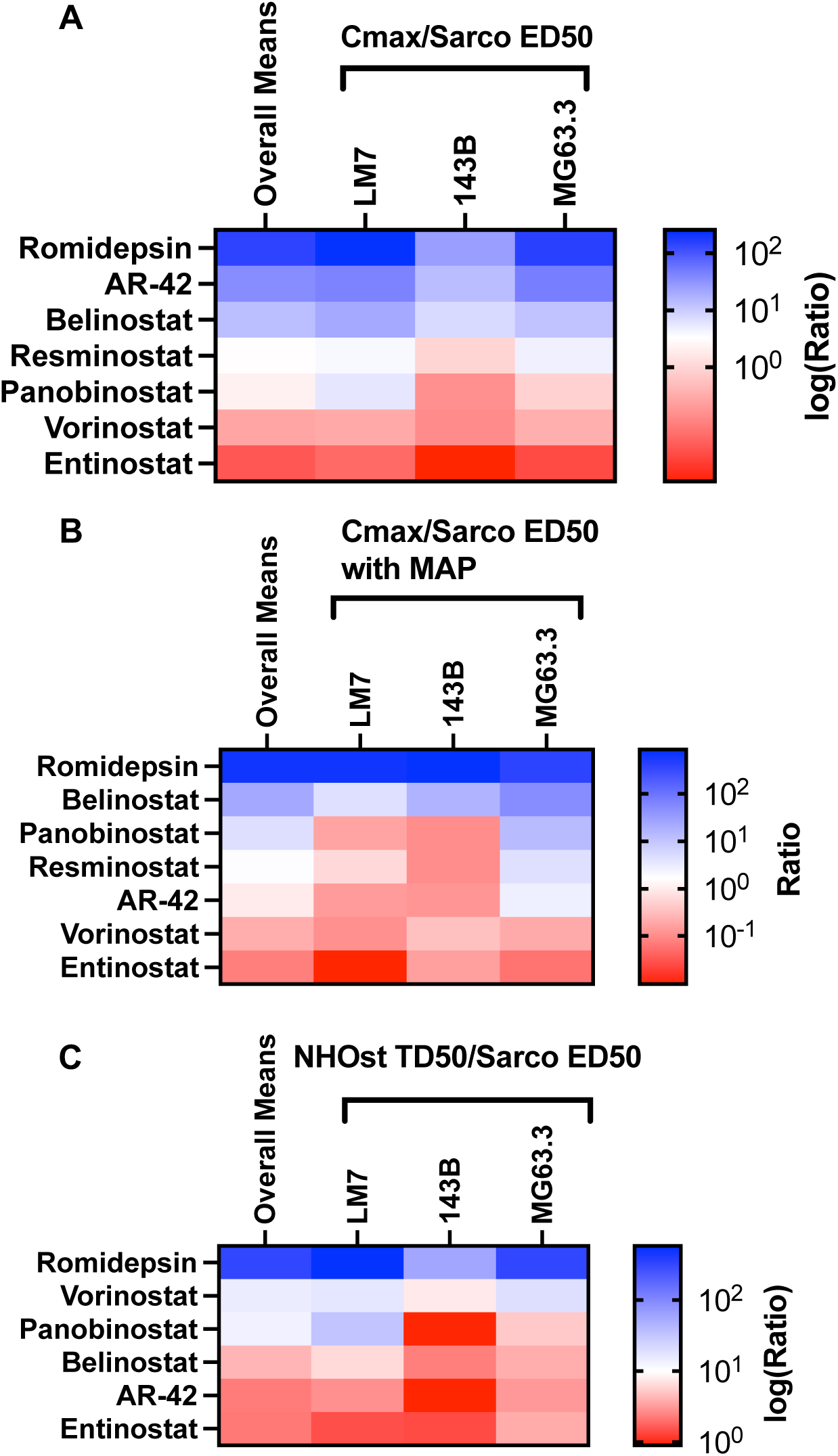
(A) Romidepsin is more promising than all other tested HDIs. Romidepsin was compared with the three other FDA-approved HDIs as well as eight that are in clinical trials. Five of the HDIs in clinical trials (Abexinostat, Givinostat, Mocetinostat, Practinostat, and Quisinostat) failed to inhibit sarcosphere viability by 50% when tested at their Cmax in preliminary studies and were therefore excluded from further study (Supplemental Figure S7). All heat maps in this Figure are sorted by the overall row means shown in the first column of each heat map. Dark blue indicates highest values (most favorable), dark red the lowest values (least favorable), and white is set at the median of the row means. **(A-B)** Ratios of Cmax/sarcosphere ED50 with and without MAP reflect potency at clinically-achievable concentrations. Cmax’s and dose responses are shown in Supplemental Table 3 and Supplemental Figure S8. ED50s were determined for sarcospheres derived from LM7, 143B, and MG63.3-GFP cell lines in N=3 independent dose response experiments with each drug. Each experiment included 3 sarcospheres per group. **(C)** Ratios of TD50/sarcosphere ED50 reflect therapeutic indices with normal human osteoblasts (NHOst). TD50s were determined in N=3 independent dose response experiments with each drug. Each experiment included at least 3 culture wells per group.

### Sarcospheres derived from a subset of human and canine osteosarcoma patients are sensitive to romidepsin

Clinical information for the human and canine osteosarcoma patients is shown in Tables 1 and 2 respectively. Human samples include primary biopsies, resections, local recurrences, and metastatic sites, which is representative of what is seen in the clinic. All canine samples were from standard of care amputations. Effects of romidepsin were measured in sarcospheres obtained from low-passage cells isolated from the human and canine osteosarcoma samples. Sarcospheres from 5 out of 18 human samples and 5 out of 10 canine samples responded to romidepsin with ED50s of 1-65 nM, which is 10- to 700-fold less than the Cmax in humans of ∼700 nM^41^ (black curves in Figure 6A-B and Tables 1-2). These groups were considered romidepsin sensitive. In contrast, sarcospheres from the other patient samples had ED50s greater than 1000 nM and were considered resistant to romidepsin (gray curves in Figure 6A-B and Tables 1-2). Romidepsin sensitivity was similar among human samples obtained from primary biopsies and post-treatment samples (primary resections, local recurrences, and metastases) as 3 out of 9 pre-treatment samples were sensitive and 2 out of 9 post treatment samples were sensitive (Table 1).

**Figure 6.**
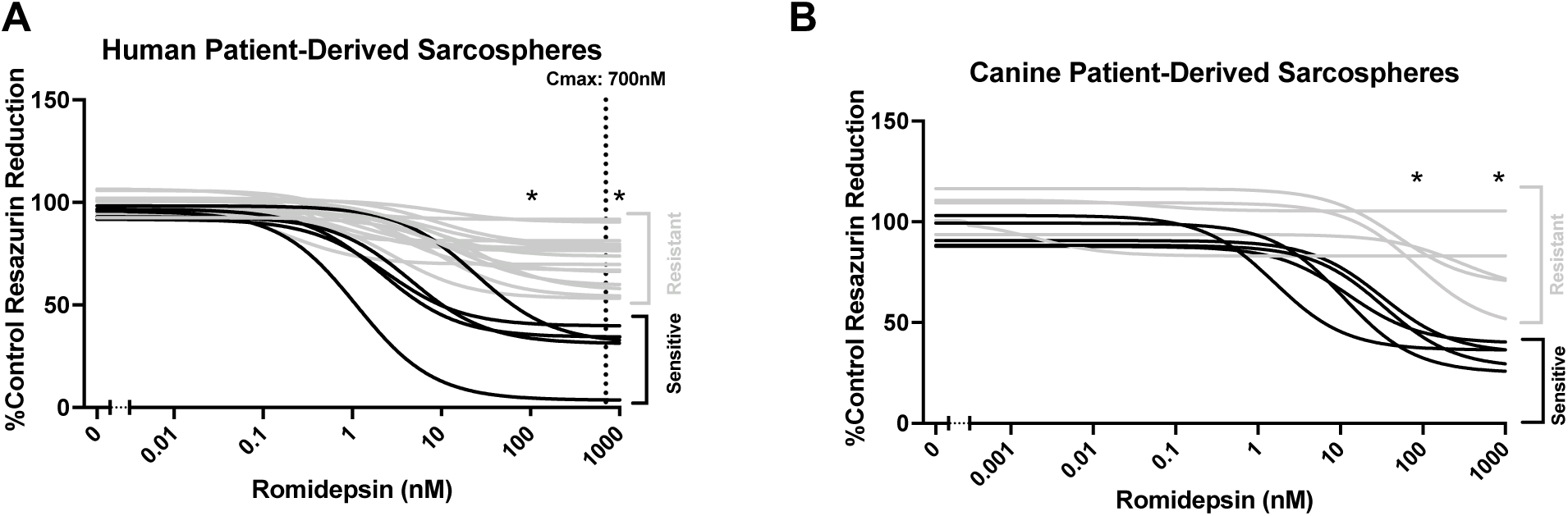
Sarcospheres from a subset of patients are sensitive to romidepsin. **(A-B)** Romidepsin dose responses on human and canine patient-derived sarcospheres. Black curves represent sensitive sarcospheres with ED50s <70 nM. Gray curves represent resistant sarcospheres, defined as ED50 >1000nM). Each dose response line represents the mean of at least 3 independent experiments with sarcospheres from an individual patient sample. Each individual experiment used 6 sarcospheres at each of the indicated doses. Asterisks represent dose groups that are different from vehicle treated group for all sensitive samples based on a one-way ANOVA with Dunnett’s multiple comparisons test (*p<0.05).

## Discussion

Long term survival in osteosarcoma has not improved over the past four decades due to therapy resistant lung metastases, indicating a need for new therapies targeting metastatic disease. We previously developed a high-throughput sarcosphere screening platform to identify promising therapeutics for metastatic osteosarcoma^23^. The current study measured effects of FDA-approved oncology drugs on the sarcospheres with or without MAP chemotherapeutics. The resultant ED50 values were compared to Cmax values from the literature and to toxicity measured on normal human osteoblasts and non-transformed human airway epithelial cells. Romidepsin, an HDI that is FDA-approved for treatment of cutaneous T-cell lymphoma^56, 57^, was the most promising candidate to repurpose for osteosarcoma among the NCI panel of 114 FDA-approved oncology drugs. The next most promising FDA-approved drugs were bortezomib and carfilzomib, the only two proteosome inhibitors in the NCI panel. Romidepsin also substantially out-performed the three other FDA-approved HDIs, and eight HDIs that are in clinical trials. Romidepsin also potently inhibited viability of sarcospheres derived at low-passage from 30-50% of human and canine patient-derived samples with ED50s substantially less than the maximally tolerated Cmax dose in humans. Our finding that sarcospheres from individual patients differ in romidepsin sensitivity suggests that sarcospheres might be useful as avatars to identify osteosarcoma patients most likely to respond clinically to romidepsin. In support of that possibility, 3D spheroid responses can predict clinical drug responses in multiple other cancers^58–61^.

The results of our unbiased, sarcosphere-based, drug screen are broadly consistent with previous osteosarcoma drug screens. For example, romidepsin, panobinostat, bortezomib, and carfilzomib were among the top hits in four independent screens with monolayers of either established osteosarcoma cell lines or PDX cell lines^62–65^. Moreover, one of the cell line screens also included drug combinations and five of the top six combinations were romidepsin with either bortezomib, carfilzomib, or the Wee1 kinase inhibitor MK1775 as well as panobinostat with carfilzomib^63^. MK1775 was not included in our screen as it is not FDA-approved. An advantage of one of the cell line monolayer screens was that dosages and exposure times were based on Cmax and half-lives of each drug in patients^63^. However, a limitation of monolayer screens is the inability to capture the complexity of an *in vivo* tumor because they do not model cell to cell interactions or biological gradients^11, 16–22^. We therefore view sarcospheres as a more clinically relevant model of micrometastatic lung metastases worth the somewhat lower throughput when compared to traditional monolayer models for drug screening.

Other investigators have also used 3D models to overcome the limitations of monolayers for osteosarcoma drug screens. For example, twenty-eight osteosarcoma patients were included in a screen based on aggregates of patient-derived cells in Matrigel (organoids)^66^. Although romidepsin was not included in that screen (A. Soragni, personal communication), five drugs were more effective at 1 uM against osteosarcomas than other sarcomas. Of those drugs, topotecan ranked 8^th^ in our initial screen at 10 uM but was not advanced to our secondary screen as its Cmax is only 6.5 nM^67^. Cabozantinib ranked 15^th^ against LM7 sarcospheres in our initial screen but had little or no effect on 143B or MG63.3 sarcospheres and everolimus had little or no effect on sarcospheres derived from all three of the cell lines. Moreover, topotecan, cabozantinib, and everolimus are already included in the National Comprehensive Cancer Network list of “Other Recommended Regimens” for osteosarcoma second line therapy. Finally, cediranib and ceralasertib were not included in our screen as they are not FDA-approved. An additional limitation of that study is that the number of drugs that could be screened was restricted by the number of cells obtained from each patient.^68^ In a screen of spheroids derived from 65 pediatric patients with various types of cancer, including seven osteosarcomas, panobinostat and bortezomib ranked 15^th^ and 19^th^ respectively of 75 drugs^69^. However, that screen did not include romidepsin or carfilzomib and the osteosarcoma results were not reported separately from the other types of cancer. Moreover, drug addition preceded spheroid formation unlike our screen where drugs were added to pre-formed sarcospheres, which is more translationally relevant as most osteosarcoma patients already have lung metastases at the time of diagnosis^3, 4^. In addition to the large drug screens described in this paragraph, a small number of FDA-approved oncology drugs were tested against a small number of osteosarcoma PDX’s^5, 6, 70–73^. However, none of those studies evaluated romidepsin, bortezomib, or carfilzomib.

In cancers with targetable driver mutations, genomic approaches can be used to select drugs for targeted therapy. However, this approach risks both false positive and false negative drug predictions in a disease without driver mutations and as genetically complex as osteosarcoma^5–8^. With further validation, emerging omics-based approaches might nonetheless be combined with patient-derived sarcospheres to identify personalized therapeutics for osteosarcoma patients. Genomics have already successfully identified pathways for targeted therapy of osteosarcoma patient-derived xenografts^5, 6, 70, 73^. Although PDXs provide a useful model to study drug effects *in vivo*, multiple roadblocks need to be overcome before they can successfully inform personalized approaches for osteosarcoma patients. Major roadblocks include that PDXs often fail to engraft and require many months to obtain results, severely reducing clinical utility and substantially increasing expense.^74–77^ For example, in a trial conducted by Champions Oncology, 24% of sarcoma PDXs failed to engraft and an additional 21% of the patients succumbed to metastatic disease prior to PDX testing results being available^78^.^78^ Similarly, in the highly immunodeficient NSG mice^79^, osteosarcoma PDX engraftment ranged from 36-50%^6, 71, 80, 81^. Predictive power is further impaired because the PDXs often do not reflect heterogeneity within the tumors^82^, are frequently implanted in non-orthotopic sites^83, 84^, and because the extended timeframe can alter drug responses by allowing genetic/epigenetic changes or infiltration of murine cells^74, 82, 85^. In contrast, every patient sample that we tested successfully formed sarcospheres and screening results can be obtained within 3 weeks, a clinically actionable timeframe.

Romidepsin substantially outperformed the other HDIs in our studies both with and without MAP chemotherapeutics. The superior effects of romidepsin compared with other HDIs are likely due to a combination of its unique mechanisms of action and its unique mechanisms of resistance^86–91^. Unique features of romidepsin compared with other HDIs include that it requires intracellular activation by reduction of the intramolecular disulfide bond^57^. Reduction of the disulfide increases romidepsin hydrophilicity likely causing increased intracellular retention^57^. Perhaps more importantly, romidepsin has slow/tight binding kinetics^52, 53^ that can lead to long-lasting effects after drug removal^51, 53, 55, 92, 93^. Romidepsin also has unique sets of HDAC targets, transcriptional co-regulators, and non-histone proteins^49, 54^, as well as unique non-transcriptional effects^51, 54, 55, 94,95^. Future studies will determine which of these unique mechanisms account for romidepsin outperforming the other HDIs in our studies.

Our results are also consistent with previous studies demonstrating that nanomolar levels of romidepsin induce cell cycle arrest and apoptosis in osteosarcoma cell lines in monolayer culture^89, 96, 97^ and that romidepsin is more potent on cancer cell lines than on non-transformed cell lines^51,98^. Romidepsin can also inhibit growth of tumors and metastases during the period of drug administration in murine osteosarcoma models. For example, growth of LM7 cell-derived lung metastases was prevented when romidepsin administration continued for the entire 21 week observation period^97^. In the same study, romidepsin had less effect on lung metastases derived from murine K7M2 osteosarcoma cells in syngeneic mice^97^. In subcutaneous xenograft models of osteosarcoma cell lines, tumors regressed when mice were followed for 1 week after discontinuation of romidepsin administration^92^. Romidepsin also inhibited growth of well-established subcutaneous xenografts in a small number of mice but did not induce a sustained response when the mice were followed for 6 weeks after romidepsin discontinuation^99^. Taken together, romidepsin appears effective in multiple murine osteosarcoma models during the period of administration but not after discontinuation. That finding is consistent with how romidepsin is used clinically as it is administered to patients with cutaneous T-cell lymphoma for as long as they continue to benefit. Future studies will determine whether romidepsin inhibits growth or metastasis of our low-passage patient-derived osteosarcoma cells in murine models.

A likely clinical application for romidepsin in osteosarcoma is combination with MAP.^24, 100^ Combination therapy can increase survival, and decrease both development of resistance and the required dosages of MAP^100^. Romidepsin had additive-to-synergistic activity with MAP in our sarcosphere experiments. Other HDIs synergize with doxorubicin or cisplatin in osteosarcoma cells^62, 64, 101–104^ and romidepsin does in other cancers.^105, 106^ Further experiments should be done to determine whether romidepsin synergizes individually with either methotrexate, doxorubicin, or cisplatin to determine whether romidepsin could potentially reduce the amount of one of the MAP chemotherapeutics administered to patients and thereby reduce lifelong toxicity^107, 108^

Together, our sarcosphere studies support further investigation of romidepsin as a potential therapeutic for osteosarcoma lung metastases. Romidepsin had higher Cmax/ED50 ratios and higher therapeutic indices (TD50/ED50 ratios) than all other drugs in the NCI panel of FDA-approved drugs, the three other FDA-approved HDIs, and another eight HDIs that are in clinical trials. Nanomolar levels of romidepsin also substantially reduced viability in sarcospheres derived from cells of 30-50% of human and canine osteosarcoma patients. That selectivity suggests sarcospheres may be useful to identify which osteosarcoma patients are most likely to benefit from romidepsin. That approach should be validated in murine models and could inform canine and human trials. Canine trials would also act as pre-clinical studies prior to human trials as osteosarcoma is highly similar in the two species and the prevalence in canines is substantially higher than in humans^24, 26, 27^.

## Acknowledgements

We thank the human patients and their families as well as the pet dogs and their families who donated tissue for this study. We also thank H. Shannon PhD (IUSM) for advice on synergy analysis as well as C. Khanna DVM, PhD (National Cancer Institute, Bethesda, MD, USA) and E.S. Kleinerman MD (MD Anderson Cancer Center, Houston, TX, USA) for providing the MG63.3-GFP and LM7 cell lines, respectively. We also thank the National Cancer Institute Developmental Therapeutics Program (NCI/DTP) for providing the panel of FDA-approved oncology drugs used in this study.

## Notes

The authors declare no potential conflicts of interest.

### Competing Interest Statement

The authors have declared no competing interest.

### Funding

- NIH: T32CA27370, R21CA209304, P30 CA082709
- Indiana University School of Medicine
- Orthopaedic Research and Education Foundation: Resident Clinician Scientist Training Grant OREF-RCSTG
- Case Western Reserve University
- Indiana Univ Simon Cancer Center
- Big Ten Cancer Research Consortium Foundation: Kenneth and Verna Mae Jessen Award
- Strides For Sarcoma
- Rally Foundation for Childhood Cancer Research
- Department of Defense: CA210123, CA230160
- The Danaher Foundation
- The Caroline Symmes Children’s Cancer Endowment
- The Tyler Trent Cancer Research Endowment

## References

(1) SEER Research Data 1973-2012 -- Surveillance, Epidemiology, and End Results (SEER) Program (www.seer.cancer.gov), National Cancer Institute, DCCPS, Surveillance Research Program, Surveillance Systems Branch, based on the November 2014 submission. (accessed.

(2) Allison, D. C.; Carney, S. C.; Ahlmann, E. R.; Hendifar, A.; Chawla, S.; Fedenko, A.; Angeles, C.; Menendez, L. R. A Meta-Analysis of Osteosarcoma Outcomes in the Modern Medical Era. Sarcoma 2012, 2012, 704872. DOI: 10.1155/2012/704872.

(3) Messerschmitt, P. J.; Garcia, R. M.; Abdul-Karim, F. W.; Greenfield, E. M.; Getty, P. J. Osteosarcoma. J Am Acad Orthop Surg 2009, 17 (8), 515–527.

(4) Jaffe, N. Historical perspective on the introduction and use of chemotherapy for the treatment of osteosarcoma. Adv Exp Med Biol 2014, 804, 1–30. DOI: 10.1007/978-3-19-04843-7_1.

(5) Pandya, P. H.; Cheng, L.; Saadatzadeh, M. R.; Bijangi-Vishehsaraei, K.; Tang, S.; Sinn, A. L.; Trowbridge, M. A.; Coy, K. L.; Bailey, B. J.; Young, C. N.;, et al. Systems Biology Approach Identifies Prognostic Signatures of Poor Overall Survival and Guides the Prioritization of Novel BET-CHK1 Combination Therapy for Osteosarcoma. Cancers (Basel) 2020, 12 (9). DOI: 10.3390/cancers12092426 From NLM PubMed-not-MEDLINE.

(6) Sayles, L. C.; Breese, M. R.; Koehne, A. L.; Leung, S. G.; Lee, A. G.; Liu, H. Y.; Spillinger, A.; Shah, A. T.; Tanasa, B.; Straessler, K.;, et al. Genome-Informed Targeted Therapy for Osteosarcoma. Cancer Discov 2019, 9 (1), 46–63. DOI: 10.1158/2159-8290.CD-17-1152 From NLM Medline.

(7) Morrow, J. J.; Khanna, C. Osteosarcoma Genetics and Epigenetics: Emerging Biology and Candidate Therapies. Crit Rev Oncog 2015, 20 (3-4), 173–197.

(8) Kovac, M.; Blattmann, C.; Ribi, S.; Smida, J.; Mueller, N. S.; Engert, F.; Castro-Giner, F.; Weischenfeldt, J.; Kovacova, M.; Krieg, A.;, et al. Exome sequencing of osteosarcoma reveals mutation signatures reminiscent of BRCA deficiency. Nat Commun 2015, 6, 8940. DOI: 10.1038/ncomms9940 From NLM.

(9) Rivera-Valentin, R. K.; Zhu, L.; Hughes, D. P. Bone Sarcomas in Pediatrics: Progress in Our Understanding of Tumor Biology and Implications for Therapy. Paediatr Drugs 2015, 17 (4), 257–271. DOI: 10.1007/s40272-015-0134-4.

(10) Gupta, S. C.; Sung, B.; Prasad, S.; Webb, L. J.; Aggarwal, B. B. Cancer drug discovery by repurposing: teaching new tricks to old dogs. Trends Pharmacol Sci 2013, 34 (9), 508–517. DOI: 10.1016/j.tips.2013.06.005.

(11) Fennema, E.; Rivron, N.; Rouwkema, J.; van Blitterswijk, C.; de Boer, J. Spheroid culture as a tool for creating 3D complex tissues. Trends Biotechnol 2013, 31 (2), 108–115. DOI: 10.1016/j.tibtech.2012.12.003 From NLM.

(12) Kimlin, L. C.; Casagrande, G.; Virador, V. M. In vitro three-dimensional (3D) models in cancer research: an update. Mol Carcinog 2013, 52 (3), 167–182. DOI: 10.1002/mc.21844 From NLM.

(13) Rimann, M.; Laternser, S.; Gvozdenovic, A.; Muff, R.; Fuchs, B.; Kelm, J. M.; Graf-Hausner, U. An in vitro osteosarcoma 3D microtissue model for drug development. J Biotechnol 2014, 189, 129–135. DOI: 10.1016/j.jbiotec.2014.09.005 From NLM.

(14) Hirschhaeuser, F.; Menne, H.; Dittfeld, C.; West, J.; Mueller-Klieser, W.; Kunz-Schughart, L. A. Multicellular tumor spheroids: an underestimated tool is catching up again. J Biotechnol 2010, 148 (1), 3–15. DOI: 10.1016/j.jbiotec.2010.01.012 From NLM.

(15) Arai, K.; Sakamoto, R.; Kubota, D.; Kondo, T. Proteomic approach toward molecular backgrounds of drug resistance of osteosarcoma cells in spheroid culture system. Proteomics 2013, 13 (15), 2351–2360. DOI: 10.1002/pmic.201300053 From NLM.

(16) Colella, G.; Fazioli, F.; Gallo, M.; De Chiara, A.; Apice, G.; Ruosi, C.; Cimmino, A.; de Nigris, F. Sarcoma Spheroids and Organoids-Promising Tools in the Era of Personalized Medicine. Int J Mol Sci 2018, 19 (2). DOI: 10.3390/ijms19020615.

(17) Friedrich, J.; Seidel, C.; Ebner, R.; Kunz-Schughart, L. A. Spheroid-based drug screen: considerations and practical approach. Nat Protoc 2009, 4 (3), 309–324. DOI: 10.1038/nprot.2008.226.

(18) Gaebler, M.; Silvestri, A.; Haybaeck, J.; Reichardt, P.; Lowery, C. D.; Stancato, L. F.; Zybarth, G.; Regenbrecht, C. R. A. Three-Dimensional Patient-Derived In Vitro Sarcoma Models: Promising Tools for Improving Clinical Tumor Management. Front Oncol 2017, 7, 203. DOI: 10.3389/fonc.2017.00203.

(19) Ivascu, A.; Kubbies, M. Rapid generation of single-tumor spheroids for high-throughput cell function and toxicity analysis. J Biomol Screen 2006, 11 (8), 922–932. DOI: 10.1177/1087057106292763.

(20) De Luca, A.; Raimondi, L.; Salamanna, F.; Carina, V.; Costa, V.; Bellavia, D.; Alessandro, R.; Fini, M.; Giavaresi, G. Relevance of 3d culture systems to study osteosarcoma environment. Journal of experimental & clinical cancer research : CR 2018, 37 (1), 2–2. DOI: 10.1186/s13046-017-0663-5 PubMed.

(21) Duval, K.; Grover, H.; Han, L. H.; Mou, Y.; Pegoraro, A. F.; Fredberg, J.; Chen, Z. Modeling Physiological Events in 2D vs. 3D Cell Culture. Physiology (Bethesda) 2017, 32 (4), 266–277. DOI: 10.1152/physiol.00036.2016 From NLM.

(22) Guan, X.; Huang, S. Advances in the application of 3D tumor models in precision oncology and drug screening. Front Bioeng Biotechnol 2022, 10, 1021966. DOI: 10.3389/fbioe.2022.1021966 From NLM.

(23) Collier, C. D.; Wirtz, E. C.; Knafler, G. J.; Morris, W. Z.; Getty, P. J.; Greenfield, E. M. Micrometastatic Drug Screening Platform Shows Heterogeneous Response to MAP Chemotherapy in Osteosarcoma Cell Lines. Clin Orthop Relat Res 2018, 476 (7), 1400–1411. DOI: 10.1007/s11999.0000000000000059.

(24) Khanna, C.; Fan, T. M.; Gorlick, R.; Helman, L. J.; Kleinerman, E. S.; Adamson, P. C.; Houghton, P. J.; Tap, W. D.; Welch, D. R.; Steeg, P. S.;, et al. Toward a drug development path that targets metastatic progression in osteosarcoma. Clin Cancer Res 2014, 20 (16), 4200–4209. DOI: 10.1158/1078-0432.Ccr-13-2574 From NLM.

(25) Koshkina, N. V.; Rao-Bindal, K.; Kleinerman, E. S. Effect of the histone deacetylase inhibitor SNDX-275 on Fas signaling in osteosarcoma cells and the feasibility of its topical application for the treatment of osteosarcoma lung metastases. Cancer 2011, 117 (15), 3457–3467. DOI: 10.1002/cncr.25884.

(26) Poon, A. C.; Matsuyama, A.; Mutsaers, A. J. Recent and current clinical trials in canine appendicular osteosarcoma. Can Vet J 2020, 61 (3), 301–308. From NLM.

(27) Simpson, S.; Dunning, M. D.; de Brot, S.; Grau-Roma, L.; Mongan, N. P.; Rutland, C. S. Comparative review of human and canine osteosarcoma: morphology, epidemiology, prognosis, treatment and genetics. Acta Vet Scand 2017, 59 (1), 71. DOI: 10.1186/s13028-017-0341-9 From NLM.

(28) Withrow, S. J.; Khanna, C. Bridging the gap between experimental animals and humans in osteosarcoma. Cancer Treat Res 2009, 152, 439–446. DOI: 10.1007/978-1-4419-0284-9_24 From NLM.

(29) Paoloni, M.; Davis, S.; Lana, S.; Withrow, S.; Sangiorgi, L.; Picci, P.; Hewitt, S.; Triche, T.; Meltzer, P.; Khanna, C. Canine tumor cross-species genomics uncovers targets linked to osteosarcoma progression. BMC Genomics 2009, 10, 625. DOI: 10.1186/1471-2164-10-625 From NLM.

(30) Selmic, L. E.; Burton, J. H.; Thamm, D. H.; Withrow, S. J.; Lana, S. E. Comparison of carboplatin and doxorubicin-based chemotherapy protocols in 470 dogs after amputation for treatment of appendicular osteosarcoma. J Vet Intern Med 2014, 28 (2), 554–563. DOI: 10.1111/jvim.12313.

(31) Ren, L.; Mendoza, A.; Zhu, J.; Briggs, J. W.; Halsey, C.; Hong, E. S.; Burkett, S. S.; Morrow, J.; Lizardo, M. M.; Osborne, T.;, et al. Characterization of the metastatic phenotype of a panel of established osteosarcoma cells. Oncotarget 2015, 6 (30), 29469–29481. DOI: 10.18632/oncotarget.5177 From NLM.

(32) Luu, H. H.; Kang, Q.; Park, J. K.; Si, W.; Luo, Q.; Jiang, W.; Yin, H.; Montag, A. G.; Simon, M. A.; Peabody, T. D.;, et al. An orthotopic model of human osteosarcoma growth and spontaneous pulmonary metastasis. Clin Exp Metastasis 2005, 22 (4), 319–329. DOI: 10.1007/s10585-005-0365-9 From NLM.

(33) Khanna, C.; Prehn, J.; Yeung, C.; Caylor, J.; Tsokos, M.; Helman, L. An orthotopic model of murine osteosarcoma with clonally related variants differing in pulmonary metastatic potential. Clin Exp Metastasis 2000, 18 (3), 261–271. DOI: 10.1023/a:1006767007547 From NLM.

(34) Duan, X.; Jia, S. F.; Zhou, Z.; Langley, R. R.; Bolontrade, M. F.; Kleinerman, E. S. Association of alphavbeta3 integrin expression with the metastatic potential and migratory and chemotactic ability of human osteosarcoma cells. Clin Exp Metastasis 2004, 21 (8), 747–753. DOI: 10.1007/s10585-005-0599-6 From NLM.

(35) Palmini, G.; Zonefrati, R.; Mavilia, C.; Aldinucci, A.; Luzi, E.; Marini, F.; Franchi, A.; Capanna, R.; Tanini, A.; Brandi, M. L. Establishment of Cancer Stem Cell Cultures from Human Conventional Osteosarcoma. J Vis Exp 2016, (116). DOI: 10.3791/53884 From NLM.

(36) Hall, M. D.; Telma, K. A.; Chang, K. E.; Lee, T. D.; Madigan, J. P.; Lloyd, J. R.; Goldlust, I. S.; Hoeschele, J. D.; Gottesman, M. M. Say no to DMSO: dimethylsulfoxide inactivates cisplatin, carboplatin, and other platinum complexes. Cancer Res 2014, 74 (14), 3913–3922. DOI: 10.1158/0008-5472.Can-14-0247 From NLM.

(37) Collier, C.; Wirtz, E.; Knafler, G.; Morris, W.; Getty, P.; Greenfield, E. Micrometastatic Drug Screening Platform Shows Heterogeneous Response to MAP Chemotherapy in Osteosarcoma Cell Lines. Clin Orthop Relat Res 2018, 476, 1400–1411. DOI: 10.1007/s11999.0000000000000059.

(38) Shoemaker, R. H. The NCI60 human tumour cell line anticancer drug screen. Nat Rev Cancer 2006, 6 (10), 813–823. DOI: 10.1038/nrc1951 From NLM.

(39) Zhang, J. H.; Chung, T. D.; Oldenburg, K. R. A Simple Statistical Parameter for Use in Evaluation and Validation of High Throughput Screening Assays. J Biomol Screen 1999, 4 (2), 67–73. DOI: 10.1177/108705719900400206 From NLM.

(40) Zheng, S.; Wang, W.; Aldahdooh, J.; Malyutina, A.; Shadbahr, T.; Tanoli, Z.; Pessia, A.; Tang, J. SynergyFinder Plus: Toward Better Interpretation and Annotation of Drug Combination Screening Datasets. Genomics Proteomics Bioinformatics 2022, 20 (3), 587–596. DOI: 10.1016/j.gpb.2022.01.004 From NLM.

(41) Liston, D. R.; Davis, M. Clinically Relevant Concentrations of Anticancer Drugs: A Guide for Nonclinical Studies. Clin Cancer Res 2017, 23 (14), 3489–3498. DOI: 10.1158/1078-0432.Ccr-16-3083 From NLM.

(42) Brunetto, A. T.; Ang, J. E.; Lal, R.; Olmos, D.; Molife, L. R.; Kristeleit, R.; Parker, A.; Casamayor, I.; Olaleye, M.; Mais, A.; et al. First-in-human, pharmacokinetic and pharmacodynamic phase I study of Resminostat, an oral histone deacetylase inhibitor, in patients with advanced solid tumors. Clin Cancer Res 2013, 19 (19), 5494–5504. DOI: 10.1158/1078-0432.Ccr-13-0735 From NLM.

(43) Sborov, D. W.; Canella, A.; Hade, E. M.; Mo, X.; Khountham, S.; Wang, J.; Ni, W.; Poi, M.; Coss, C.; Liu, Z.;, et al. A phase 1 trial of the HDAC inhibitor AR-42 in patients with multiple myeloma and T- and B-cell lymphomas. Leuk Lymphoma 2017, 58 (10), 2310–2318. DOI: 10.1080/10428194.2017.1298751 From NLM.

(44) Pili, R.; Salumbides, B.; Zhao, M.; Altiok, S.; Qian, D.; Zwiebel, J.; Carducci, M. A.; Rudek, M. A. Phase I study of the histone deacetylase inhibitor entinostat in combination with 13-cis retinoic acid in patients with solid tumours. Br J Cancer 2012, 106 (1), 77–84. DOI: 10.1038/bjc.2011.527 From NLM.

(45) Grohar, P. J.; Glod, J.; Peer, C. J.; Sissung, T. M.; Arnaldez, F. I.; Long, L.; Figg, W. D.; Whitcomb, P.; Helman, L. J.; Widemann, B. C. A phase I/II trial and pharmacokinetic study of mithramycin in children and adults with refractory Ewing sarcoma and EWS-FLI1 fusion transcript. Cancer Chemother Pharmacol 2017, 80 (3), 645–652. DOI: 10.1007/s00280-017-3382-x From NLM.

(46) Ianevski, A.; Giri, A. K.; Aittokallio, T. SynergyFinder 3.0: an interactive analysis and consensus interpretation of multi-drug synergies across multiple samples. Nucleic Acids Research 2022, 50 (W1), W739–W743. DOI: 10.1093/nar/gkac382 (acccessed 7/18/2024).

(47) Gordon, N.; Kleinerman, E. S. Aerosol therapy for the treatment of osteosarcoma lung metastases: targeting the Fas/FasL pathway and rationale for the use of gemcitabine. J Aerosol Med Pulm Drug Deliv 2010, 23 (4), 189–196. DOI: 10.1089/jamp.2009.0812 From NLM.

(48) Rodriguez, C. O., Jr.; Crabbs, T. A.; Wilson, D. W.; Cannan, V. A.; Skorupski, K. A.; Gordon, N.; Koshkina, N.; Kleinerman, E.; Anderson, P. M. Aerosol gemcitabine: preclinical safety and in vivo antitumor activity in osteosarcoma-bearing dogs. J Aerosol Med Pulm Drug Deliv 2010, 23 (4), 197–206. DOI: 10.1089/jamp.2009.0773.

(49) Bantscheff, M.; Hopf, C.; Savitski, M. M.; Dittmann, A.; Grandi, P.; Michon, A. M.; Schlegl, J.; Abraham, Y.; Becher, I.; Bergamini, G.;, et al. Chemoproteomics profiling of HDAC inhibitors reveals selective targeting of HDAC complexes. Nat Biotechnol 2011, 29 (3), 255–265. DOI: 10.1038/nbt.1759 From NLM.

(50) Fraczek, J.; Vanhaecke, T.; Rogiers, V. Toxicological and metabolic considerations for histone deacetylase inhibitors. Expert Opin Drug Metab Toxicol 2013, 9 (4), 441–457. DOI: 10.1517/17425255.2013.754011 From NLM.

(51) Safari, M.; Litman, T.; Robey, R. W.; Aguilera, A.; Chakraborty, A. R.; Reinhold, W. C.; Basseville, A.; Petrukhin, L.; Scotto, L.; O’Connor, O. A.;, et al. R-Loop-Mediated ssDNA Breaks Accumulate Following Short-Term Exposure to the HDAC Inhibitor Romidepsin. Mol Cancer Res 2021, 19 (8), 1361–1374. DOI: 10.1158/1541-7786.Mcr-20-0833 From NLM.

(52) Kitir, B.; Maolanon, A. R.; Ohm, R. G.; Colaço, A. R.; Fristrup, P.; Madsen, A. S.; Olsen, C. A. Chemical Editing of Macrocyclic Natural Products and Kinetic Profiling Reveal Slow, Tight-Binding Histone Deacetylase Inhibitors with Picomolar Affinities. Biochemistry 2017, 56 (38), 5134–5146. DOI: 10.1021/acs.biochem.7b00725 From NLM.

(53) Robers, M. B.; Dart, M. L.; Woodroofe, C. C.; Zimprich, C. A.; Kirkland, T. A.; Machleidt, T.; Kupcho, K. R.; Levin, S.; Hartnett, J. R.; Zimmerman, K.;, et al. Target engagement and drug residence time can be observed in living cells with BRET. Nat Commun 2015, 6, 10091. DOI: 10.1038/ncomms10091 From NLM.

(54) Lechner, S.; Malgapo, M. I. P.; Grätz, C.; Steimbach, R. R.; Baron, A.; Rüther, P.; Nadal, S.; Stumpf, C.; Loos, C.; Ku, X.;, et al. Target deconvolution of HDAC pharmacopoeia reveals MBLAC2 as common off-target. Nat Chem Biol 2022, 18 (8), 812–820. DOI: 10.1038/s41589-022-01015-5 From NLM.

(55) Luchenko, V. L.; Litman, T.; Chakraborty, A. R.; Heffner, A.; Devor, C.; Wilkerson, J.; Stein, W.; Robey, R. W.; Bangiolo, L.; Levens, D.;, et al. Histone deacetylase inhibitor-mediated cell death is distinct from its global effect on chromatin. Mol Oncol 2014, 8 (8), 1379–1392. DOI: 10.1016/j.molonc.2014.05.001 From NLM.

(56) Grant, C.; Rahman, F.; Piekarz, R.; Peer, C.; Frye, R.; Robey, R. W.; Gardner, E. R.; Figg, W. D.; Bates, S. E. Romidepsin: a new therapy for cutaneous T-cell lymphoma and a potential therapy for solid tumors. Expert Rev Anticancer Ther 2010, 10 (7), 997–1008. DOI: 10.1586/era.10.88 From NLM.

(57) Furumai, R.; Matsuyama, A.; Kobashi, N.; Lee, K. H.; Nishiyama, M.; Nakajima, H.; Tanaka, A.; Komatsu, Y.; Nishino, N.; Yoshida, M.;, et al. FK228 (depsipeptide) as a natural prodrug that inhibits class I histone deacetylases. Cancer Res 2002, 62 (17), 4916–4921. From NLM.

(58) Halfter, K.; Hoffmann, O.; Ditsch, N.; Ahne, M.; Arnold, F.; Paepke, S.; Grab, D.; Bauerfeind, I.; Mayer, B. Testing chemotherapy efficacy in HER2 negative breast cancer using patient-derived spheroids. J Transl Med 2016, 14 (1), 112. DOI: 10.1186/s12967-016-0855-3.

(59) Shuford, S.; Wilhelm, C.; Rayner, M.; Elrod, A.; Millard, M.; Mattingly, C.; Lotstein, A.; Smith, A. M.; Guo, Q. J.; O’Donnell, L.;, et al. Prospective Validation of an Ex Vivo, Patient-Derived 3D Spheroid Model for Response Predictions in Newly Diagnosed Ovarian Cancer. Sci Rep 2019, 9 (1), 11153. DOI: 10.1038/s41598-019-47578-7.

(60) Shuford, S.; Lipinski, L.; Abad, A.; Smith, A. M.; Rayner, M.; O’Donnell, L.; Stuart, J.; Mechtler, L. L.; Fabiano, A. J.; Edenfield, J.;, et al. Prospective prediction of clinical drug response in high-grade gliomas using an ex vivo 3D cell culture assay. Neurooncol Adv 2021, 3 (1), vdab065. DOI: 10.1093/noajnl/vdab065.

(61) Åkerlund, E.; Gudoityte, G.; Moussaud-Lamodière, E.; Lind, O.; Bwanika, H. C.; Lehti, K.; Salehi, S.; Carlson, J.; Wallin, E.; Fernebro, J.;, et al. The drug efficacy testing in 3D cultures platform identifies effective drugs for ovarian cancer patients. NPJ Precis Oncol 2023, 7 (1), 111. DOI: 10.1038/s41698-023-00463-z From NLM.

(62) Loh, A. H. P.; Stewart, E.; Bradley, C. L.; Chen, X.; Daryani, V.; Stewart, C. F.; Calabrese, C.; Funk, A.; Miller, G.; Karlstrom, A.;, et al. Combinatorial screening using orthotopic patient derived xenograft-expanded early phase cultures of osteosarcoma identify novel therapeutic drug combinations. Cancer Lett 2019, 442, 262–270. DOI: 10.1016/j.canlet.2018.10.033 From NLM.

(63) Yu, D.; Kahen, E.; Cubitt, C. L.; McGuire, J.; Kreahling, J.; Lee, J.; Altiok, S.; Lynch, C. C.; Sullivan, D. M.; Reed, D. R. Identification of Synergistic, Clinically Achievable, Combination Therapies for Osteosarcoma. Sci Rep 2015, 5, 16991. DOI: 10.1038/srep16991 From NLM.

(64) Zhang, W.; Qi, L.; Liu, Z.; He, S.; Wang, C. Z.; Wu, Y.; Han, L.; Liu, Z.; Fu, Z.; Tu, C.;, et al. Integrated multiomic analysis and high-throughput screening reveal potential gene targets and synergetic drug combinations for osteosarcoma therapy. MedComm (2020) 2023, 4 (4), e317. DOI: 10.1002/mco2.317 From NLM.

(65) Khalid, U.; Simovic, M.; Hammann, L. A.; Iskar, M.; Wong, J. K. L.; Kumar, R.; Jugold, M.; Sill, M.; Bolkestein, M.; Kolb, T.;, et al. A synergistic interaction between HDAC- and PARP inhibitors in childhood tumors with chromothripsis. Int J Cancer 2022, 151 (4), 590–606. DOI: 10.1002/ijc.34027 From NLM.

(66) Al Shihabi, A.; Tebon, P. J.; Nguyen, H. T. L.; Chantharasamee, J.; Sartini, S.; Davarifar, A.; Jensen, A. Y.; Diaz-Infante, M.; Cox, H.; Gonzalez, A. E.;, et al. The landscape of drug sensitivity and resistance in sarcoma. Cell Stem Cell 2024, 31 (10), 1524–1542.e1524. DOI: 10.1016/j.stem.2024.08.010 From NLM.

(67) Jamaladdin, N.; Sigaud, R.; Kocher, D.; Kolodziejczak, A. S.; Nonnenbroich, L. F.; Ecker, J.; Usta, D.; Benzel, J.; Peterziel, H.; Pajtler, K. W.;, et al. Key Pharmacokinetic Parameters of 74 Pediatric Anticancer Drugs Providing Assistance in Preclinical Studies. Clin Pharmacol Ther 2023, 114 (4), 904–913. DOI: 10.1002/cpt.3002 From NLM.

(68) Al Shihabi, A.; Tebon, P. J.; Nguyen, H. T. L.; Chantharasamee, J.; Sartini, S.; Davarifar, A.; Jensen, A. Y.; Diaz-Infante, M.; Cox, H.; Gonzalez, A. E.; et al. The landscape of drug sensitivity and resistance in sarcoma. bioRxiv 2023, 2023.2005.2025.542375. DOI: 10.1101/2023.05.25.542375.

(69) Peterziel, H.; Jamaladdin, N.; ElHarouni, D.; Gerloff, X. F.; Herter, S.; Fiesel, P.; Berker, Y.; Blattner-Johnson, M.; Schramm, K.; Jones, B. C.;, et al. Drug sensitivity profiling of 3D tumor tissue cultures in the pediatric precision oncology program INFORM. NPJ Precis Oncol 2022, 6 (1), 94. DOI: 10.1038/s41698-022-00335-y From NLM.

(70) Schott, C. R.; Koehne, A. L.; Sayles, L. C.; Young, E. P.; Luck, C.; Yu, K.; Lee, A. G.; Breese, M. R.; Leung, S. G.; Xu, H.;, et al. Osteosarcoma PDX-Derived Cell Line Models for Preclinical Drug Evaluation Demonstrate Metastasis Inhibition by Dinaciclib through a Genome-Targeted Approach. Clin Cancer Res 2024, 30 (4), 849–864. DOI: 10.1158/1078-0432.Ccr-23-0873 From NLM.

(71) Blankenship, K.; Chang, T.-C.; Fan, Y.; Gordon, B.; Wright, W. C.; Kieffer, M.; Karlstrom, A.; Jeon, J.; Patel, A. G.; Dapper, J.;, et al. Preclinical Pediatric Molecular Analysis for Therapy Choice (MATCH). bioRxiv 2024, 2024.2007.2022.604648. DOI: 10.1101/2024.07.22.604648.

(72) Castillo-Ecija, H.; Pascual-Pasto, G.; Perez-Jaume, S.; Resa-Pares, C.; Vila-Ubach, M.; Monterrubio, C.; Jimenez-Cabaco, A.; Baulenas-Farres, M.; Muñoz-Aznar, O.; Salvador, N.;, et al. Prognostic value of patient-derived xenograft engraftment in pediatric sarcomas. J Pathol Clin Res 2021, 7 (4), 338–349. DOI: 10.1002/cjp2.210 From NLM.

(73) Pandya, P. H.; Jannu, A. J.; Bijangi-Vishehsaraei, K.; Dobrota, E.; Bailey, B. J.; Barghi, F.; Shannon, H. E.; Riyahi, N.; Damayanti, N. P.; Young, C.;, et al. Integrative Multi-OMICs Identifies Therapeutic Response Biomarkers and Confirms Fidelity of Clinically Annotated, Serially Passaged Patient-Derived Xenografts Established from Primary and Metastatic Pediatric and AYA Solid Tumors. Cancers (Basel) 2022, 15 (1). DOI: 10.3390/cancers15010259 From NLM.

(74) Friedman, A. A.; Letai, A.; Fisher, D. E.; Flaherty, K. T. Precision medicine for cancer with next-generation functional diagnostics. Nat Rev Cancer 2015, 15 (12), 747–756. DOI: 10.1038/nrc4015.

(75) Halfter, K.; Mayer, B. Bringing 3D tumor models to the clinic -predictive value for personalized medicine. Biotechnol J 2017, 12 (2), 1600295. DOI: 10.1002/biot.201600295.

(76) Hingorani, P.; Janeway, K.; Crompton, B. D.; Kadoch, C.; Mackall, C. L.; Khan, J.; Shern, J. F.; Schiffman, J.; Mirabello, L.; Savage, S. A.;, et al. Current state of pediatric sarcoma biology and opportunities for future discovery: A report from the sarcoma translational research workshop. Cancer Genet 2016, 209 (5), 182–194. DOI: 10.1016/j.cancergen.2016.03.004.

(77) Izumchenko, E.; Paz, K.; Ciznadija, D.; Sloma, I.; Katz, A.; Vasquez-Dunddel, D.; Ben-Zvi, I.; Stebbing, J.; McGuire, W.; Harris, W.;, et al. Patient-derived xenografts effectively capture responses to oncology therapy in a heterogeneous cohort of patients with solid tumors. Ann Oncol 2017, 28 (10), 2595–2605. DOI: 10.1093/annonc/mdx416.

(78) Stebbing, J.; Paz, K.; Schwartz, G. K.; Wexler, L. H.; Maki, R.; Pollock, R. E.; Morris, R.; Cohen, R.; Shankar, A.; Blackman, G.;, et al. Patient-derived xenografts for individualized care in advanced sarcoma. Cancer 2014, 120 (13), 2006–2015. DOI: 10.1002/cncr.28696 From NLM.

(79) Okada, S.; Vaeteewoottacharn, K.; Kariya, R. Application of Highly Immunocompromised Mice for the Establishment of Patient-Derived Xenograft (PDX) Models. Cells 2019, 8 (8). DOI: 10.3390/cells8080889 From NLM.

(80) Stewart, E.; Federico, S. M.; Chen, X.; Shelat, A. A.; Bradley, C.; Gordon, B.; Karlstrom, A.; Twarog, N. R.; Clay, M. R.; Bahrami, A.;, et al. Orthotopic patient-derived xenografts of paediatric solid tumours. Nature 2017, 549 (7670), 96–100. DOI: 10.1038/nature23647 From NLM.

(81) Nanni, P.; Landuzzi, L.; Manara, M. C.; Righi, A.; Nicoletti, G.; Cristalli, C.; Pasello, M.; Parra, A.; Carrabotta, M.; Ferracin, M.;, et al. Bone sarcoma patient-derived xenografts are faithful and stable preclinical models for molecular and therapeutic investigations. Sci Rep 2019, 9 (1), 12174. DOI: 10.1038/s41598-019-48634-y From NLM.

(82) Yoshida, G. J. Applications of patient-derived tumor xenograft models and tumor organoids. J Hematol Oncol 2020, 13 (1), 4. DOI: 10.1186/s13045-019-0829-z From NLM.

(83) Hidalgo, M.; Amant, F.; Biankin, A. V.; Budinská, E.; Byrne, A. T.; Caldas, C.; Clarke, R. B.; de Jong, S.; Jonkers, J.; Mælandsmo, G. M.;, et al. Patient-derived xenograft models: an emerging platform for translational cancer research. Cancer Discov 2014, 4 (9), 998–1013. DOI: 10.1158/2159-8290.Cd-14-0001 From NLM.

(84) Hiroshima, Y.; Maawy, A.; Zhang, Y.; Zhang, N.; Murakami, T.; Chishima, T.; Tanaka, K.; Ichikawa, Y.; Bouvet, M.; Endo, I.;, et al. Patient-derived mouse models of cancer need to be orthotopic in order to evaluate targeted anti-metastatic therapy. Oncotarget 2016, 7 (44), 71696–71702. DOI: 10.18632/oncotarget.12322 From NLM.

(85) Ben-David, U.; Ha, G.; Tseng, Y. Y.; Greenwald, N. F.; Oh, C.; Shih, J.; McFarland, J. M.; Wong, B.; Boehm, J. S.; Beroukhim, R.;, et al. Patient-derived xenografts undergo mouse-specific tumor evolution. Nat Genet 2017. DOI: 10.1038/ng.3967.

(86) Kanno, S.; Maeda, N.; Tomizawa, A.; Yomogida, S.; Katoh, T.; Ishikawa, M. Characterization of cells resistant to the potent histone deacetylase inhibitor spiruchostatin B (SP-B) and effect of overexpressed p21waf1/cip1 on the SP-B resistance or susceptibility of human leukemia cells. Int J Oncol 2012, 41 (3), 862–868. DOI: 10.3892/ijo.2012.1507 From NLM Medline.

(87) Robey, R. W.; Fitzsimmons, C. M.; Guiblet, W. M.; Frye, W. J. E.; González Dalmasy, J. M.; Wang, L.; Russell, D. A.; Huff, L. M.; Perciaccante, A. J.; Ali-Rahmani, F.;, et al. The Methyltransferases METTL7A and METTL7B Confer Resistance to Thiol-Based Histone Deacetylase Inhibitors. Mol Cancer Ther 2024, 23 (4), 464–477. DOI: 10.1158/1535-7163.Mct-23-0144 From NLM.

(88) Xiao, J. J.; Huang, Y.; Dai, Z.; Sadée, W.; Chen, J.; Liu, S.; Marcucci, G.; Byrd, J.; Covey, J. M.; Wright, J.; et al. Chemoresistance to depsipeptide FK228 [(E)-(1S,4S,10S,21R)-7-[(Z)-ethylidene]-4,21-diisopropyl-2-oxa-12,13-dithia-5,8,20,23-tetraazabicyclo[8,7,6]-tricos-16-ene-3,6,9,22-pentanone] is mediated by reversible MDR1 induction in human cancer cell lines. J Pharmacol Exp Ther 2005, 314 (1), 467–475. DOI: 10.1124/jpet.105.083956 From NLM.

(89) Matsubara, H.; Watanabe, M.; Imai, T.; Yui, Y.; Mizushima, Y.; Hiraumi, Y.; Kamitsuji, Y.; Watanabe, K.; Nishijo, K.; Toguchida, J.; et al. Involvement of extracellular signal-regulated kinase activation in human osteosarcoma cell resistance to the histone deacetylase inhibitor FK228 [(1S,4S,7Z,10S,16E,21R)-7-ethylidene-4,21-bis(propan-2-yl)-2-oxa-12,13-dithia-5,8,20,23-tetraazabicyclo[8.7.6]tricos-16-ene-3,6,9,19,22-pentone]. J Pharmacol Exp Ther 2009, 328 (3), 839–848. DOI: 10.1124/jpet.108.147462 From NLM.

(90) Chen, G.; Li, A.; Zhao, M.; Gao, Y.; Zhou, T.; Xu, Y.; Du, Z.; Zhang, X.; Yu, X. Proteomic analysis identifies protein targets responsible for depsipeptide sensitivity in tumor cells. J Proteome Res 2008, 7 (7), 2733–2742. DOI: 10.1021/pr7008753 From NLM.

(91) Okada, T.; Tanaka, K.; Nakatani, F.; Sakimura, R.; Matsunobu, T.; Li, X.; Hanada, M.; Nakamura, T.; Oda, Y.; Tsuneyoshi, M.;, et al. Involvement of P-glycoprotein and MRP1 in resistance to cyclic tetrapeptide subfamily of histone deacetylase inhibitors in the drug-resistant osteosarcoma and Ewing’s sarcoma cells. Int J Cancer 2006, 118 (1), 90–97. DOI: 10.1002/ijc.21297 From NLM.

(92) Imai, T.; Adachi, S.; Nishijo, K.; Ohgushi, M.; Okada, M.; Yasumi, T.; Watanabe, K.; Nishikomori, R.; Nakayama, T.; Yonehara, S.;, et al. FR901228 induces tumor regression associated with induction of Fas ligand and activation of Fas signaling in human osteosarcoma cells. Oncogene 2003, 22 (58), 9231–9242. DOI: 10.1038/sj.onc.1207184 From NLM.

(93) Lauffer, B. E.; Mintzer, R.; Fong, R.; Mukund, S.; Tam, C.; Zilberleyb, I.; Flicke, B.; Ritscher, A.; Fedorowicz, G.; Vallero, R.;, et al. Histone deacetylase (HDAC) inhibitor kinetic rate constants correlate with cellular histone acetylation but not transcription and cell viability. J Biol Chem 2013, 288 (37), 26926–26943. DOI: 10.1074/jbc.M113.490706 From NLM.

(94) Lutz, L.; Fitzner, I. C.; Ahrens, T.; Geißler, A. L.; Makowiec, F.; Hopt, U. T.; Bogatyreva, L.; Hauschke, D.; Werner, M.; Lassmann, S. Histone modifiers and marks define heterogeneous groups of colorectal carcinomas and affect responses to HDAC inhibitors in vitro. Am J Cancer Res 2016, 6 (3), 664–676. From NLM.

(95) Chang, J.; Varghese, D. S.; Gillam, M. C.; Peyton, M.; Modi, B.; Schiltz, R. L.; Girard, L.; Martinez, E. D. Differential response of cancer cells to HDAC inhibitors trichostatin A and depsipeptide. Br J Cancer 2012, 106 (1), 116–125. DOI: 10.1038/bjc.2011.532 From NLM.

(96) Watanabe, K.; Okamoto, K.; Yonehara, S. Sensitization of osteosarcoma cells to death receptor-mediated apoptosis by HDAC inhibitors through downregulation of cellular FLIP. Cell Death Differ 2005, 12 (1), 10–18. DOI: 10.1038/sj.cdd.4401507 From NLM.

(97) McGuire, J. J.; Nerlakanti, N.; Lo, C. H.; Tauro, M.; Utset-Ward, T. J.; Reed, D. R.; Lynch, C. C. Histone deacetylase inhibition prevents the growth of primary and metastatic osteosarcoma. Int J Cancer 2020, 147 (10), 2811–2823. DOI: 10.1002/ijc.33046 From NLM.

(98) Paillas, S.; Then, C. K.; Kilgas, S.; Ruan, J. L.; Thompson, J.; Elliott, A.; Smart, S.; Kiltie, A. E. The Histone Deacetylase Inhibitor Romidepsin Spares Normal Tissues While Acting as an Effective Radiosensitizer in Bladder Tumors in Vivo. Int J Radiat Oncol Biol Phys 2020, 107 (1), 212–221. DOI: 10.1016/j.ijrobp.2020.01.015 From NLM Medline.

(99) Graham, C.; Tucker, C.; Creech, J.; Favours, E.; Billups, C. A.; Liu, T.; Fouladi, M.; Freeman, B. B., 3rd; Stewart, C. F.; Houghton, P. J. Evaluation of the antitumor efficacy, pharmacokinetics, and pharmacodynamics of the histone deacetylase inhibitor depsipeptide in childhood cancer models in vivo. Clin Cancer Res 2006, 12 (1), 223–234. DOI: 10.1158/1078-0432.Ccr-05-1225 From NLM.

(100) Collier, C. D.; Getty, P. J.; Greenfield, E. M. Targeting the Cancer Epigenome with Histone Deacetylase Inhibitors in Osteosarcoma. Adv Exp Med Biol 2020, 1258, 55–75. DOI: 10.1007/978-3-030-43085-6_4 From NLM.

(101) Wittenburg, L. A.; Bisson, L.; Rose, B. J.; Korch, C.; Thamm, D. H. The histone deacetylase inhibitor valproic acid sensitizes human and canine osteosarcoma to doxorubicin. Cancer Chemother Pharmacol 2011, 67 (1), 83–92. DOI: 10.1007/s00280-010-1287-z.

(102) Yang, C.; Choy, E.; Hornicek, F. J.; Wood, K. B.; Schwab, J. H.; Liu, X.; Mankin, H.; Duan, Z. Histone deacetylase inhibitor (HDACI) PCI-24781 potentiates cytotoxic effects of doxorubicin in bone sarcoma cells. Cancer Chemother Pharmacol 2011, 67 (2), 439–446. DOI: 10.1007/s00280-010-1344-7.

(103) Pettke, A.; Hotfilder, M.; Clemens, D.; Klco-Brosius, S.; Schaefer, C.; Potratz, J.; Dirksen, U. Suberanilohydroxamic acid (vorinostat) synergistically enhances the cytotoxicity of doxorubicin and cisplatin in osteosarcoma cell lines. Anticancer Drugs 2016, 27 (10), 1001–1010. DOI: 10.1097/CAD.0000000000000418.

(104) Murahari, S.; Jalkanen, A. L.; Kulp, S. K.; Chen, C. S.; Modiano, J. F.; London, C. A.; Kisseberth, W. C. Sensitivity of osteosarcoma cells to HDAC inhibitor AR-42 mediated apoptosis. BMC Cancer 2017, 17 (1), 67. DOI: 10.1186/s12885-017-3046-6.

(105) Luchenko, V. L.; Salcido, C. D.; Zhang, Y.; Agama, K.; Komlodi-Pasztor, E.; Murphy, R. F.; Giaccone, G.; Pommier, Y.; Bates, S. E.; Varticovski, L. Schedule-dependent synergy of histone deacetylase inhibitors with DNA damaging agents in small cell lung cancer. Cell Cycle 2011, 10 (18), 3119–3128. DOI: 10.4161/cc.10.18.17190.

(106) Stiborova, M.; Eckschlager, T.; Poljakova, J.; Hrabeta, J.; Adam, V.; Kizek, R.; Frei, E. The synergistic effects of DNA-targeted chemotherapeutics and histone deacetylase inhibitors as therapeutic strategies for cancer treatment. Curr Med Chem 2012, 19 (25), 4218–4238.

(107) Bhagat, A.; Kleinerman, E. S. Anthracycline-Induced Cardiotoxicity: Causes, Mechanisms, and Prevention. Adv Exp Med Biol 2020, 1257, 181–192. DOI: 10.1007/978-3-030-43032-0_15 From NLM.

(108) Janeway, K. A.; Grier, H. E. Sequelae of osteosarcoma medical therapy: a review of rare acute toxicities and late effects. Lancet Oncol 2010, 11 (7), 670–678. DOI: 10.1016/s1470-2045(10)70062-0 From NLM.

