## Supplemental Data for "Repurposing Romidepsin for Osteosarcoma: Screening FDA-Approved Oncology Drugs with Three-Dimensional Osteosarcoma Spheroids"

| Supplemental Table 1. Suppliers and Storage Conditions for 13 Drugs in Secondary Screen | | |
|---|---|---|
| **Drug** | **Supplier** | **Storage** |
| Bortezomib / PS-341 | Boston Biochem / R&D | -20C |
| Romidepsin / FK-228 | Tocris / R&D | -20C |
| Plicamycin / NSC-24559 | DTP / Sigma | Dark, -20C |
| Mitomycin C / NSC-26980 | DTP/ APAC Pharm | Dark, 4C |
| Teniposide / NSC-122819 | DTP / Sigma | Dark, -20C |
| Mitoxantrone / NSC-279836 | DTP | -20C |
| Vorinostat / NSC-701852 | DTP / Sigma | -20C |
| Afatinib / NSC-750691 | DTP | -20C |
| Crizotinib / NSC-756645 | DTP | Desiccate, -20C |
| Carfilzomib / NSC-758252 | DTP | Desiccate, dark, -20C |
| Omacetaxine / NSC-758253 | DTP | -20C |
| Ponatinib / NSC-758487 | DTP | -20C |
| Vandetanib / NSC-760766 | DTP/Ontario Chem | -20C |

DTP: Developmental Therapeutics Program (NCI)

| Supplemental Table 2. Concentrations of MAP Chemotherapeutics Used (nM) | | | |
|---|---|---|---|
| **Initial Screen: To obtain GIC20 with methotrexate, doxorubicin, and cisplatin in combination** | | | |
| **Drug** | **LM7** | **143B** | **MG63.3** |
| Methotrexate | 61,091 | 4.83 | 8,210 |
| Doxorubicin | 27.9 | 48.3 | 58 |
| Cisplatin | 104.7 | 108.7 | 154.7 |
| **Secondary Screen: To obtain ED50 with methotrexate, doxorubicin, and cisplatin in combination** | | | |
| **Drug** | **LM7** | **143B** | **MG63.3** |
| Methotrexate | 618,963 | 30 | 67,655 |
| Doxorubicin | 283 | 301 | 478 |
| Cisplatin | 1,061 | 678 | 1,274 |

**Supplemental Table 3. Cmax and Sarcosphere ED50s for 13 Hits in Secondary Screen and Histone Deacetylase Inhibitors**

| Secondary Screen | Cmax[41] | ED50 LM7 | ED50 143B | ED50 MG63.3-GFP | ED50 w/ MAP LM7 | ED50 w/ MAP 143B | ED50 w/MAP MG63.3-GFP |
|---|---|---|---|---|---|---|---|
| Afatinib | 52 | 2590 | 3959 | 2251 | 6918 | 4174 | 8817 |
| Bortezomib | 312 | 322 | 3.5 | 2.3 | 1477 | 7.4 | 3.7 |
| Carfilzomib | 5880 | 528 | 39.5 | 18 | >10,000 | 199 | 35.4 |
| Crizotinib | 514 | 2520 | 2609 | 2052 | 2824 | 2592 | 4122 |
| Mitomycin C | 2180 | 1375 | 899 | 1375 | 2282 | 913 | 3287 |
| Mixoxantrone | 715 | 240 | 113 | 253 | 944 | 626 | 1032 |
| Omacetizine | 46 | 441 | 63.1 | 68.9 | 212 | 180 | 105 |
| Plicamycin | 18[45] | 93.3 | 23.6 | 42.3 | >10,000 | 5.1 | 184 |
| Ponatinib | 137 | 630 | 1167 | 1237 | 405 | 1135 | 2372 |
| Romidepsin | 697 | 2.76 | 28.1 | 4.81 | 1.11 | 0.81 | 1.80 |
| Teniposide | 23,100 | >10,000 | 267 | 1561 | >10,000 | 281 | 2544 |
| Vandetanib | 2160 | 3470 | 5305 | 4514 | 5469 | 2681 | 5164 |
| Vorinostat | 1200 | 4167 | 8944 | 3613 | 1359 | >10,000 | 3526 |
| **HDIs** | | | | | | | |
| Romidepsin | 697 | 2.8 | 28.4 | 4.9 | 1.1 | 0.8 | 1.8 |
| AR-42 | 1500[43] | 1310 | 4110 | 1190 | >10,000 | >10,000 | 563 |
| Belinostat | 134,000 | 5079 | >10,000 | 8336 | >10,000 | 8221 | 2779 |
| Panobinostat | 82 | 8.7 | 320 | 16.1 | 477 | 781 | 6.3 |
| Vorinostat | 1200 | 4167 | 8944 | 3614 | 1359 | >10,000 | 3526 |
| Resminostat | 14220[42] | 3160 | >10,000 | 2650 | >10,000 | >10,000 | 3260 |
| Entinostat | 289[44] | 6165 | >10,000 | 7060 | >10,000 | 1846 | 4843 |

All units for values listed are nM

All Cmax values except for Plicamycin, AR-42, Entinostat, Resminostat are from reference 41. References for those Cmax values are indicated in the table.

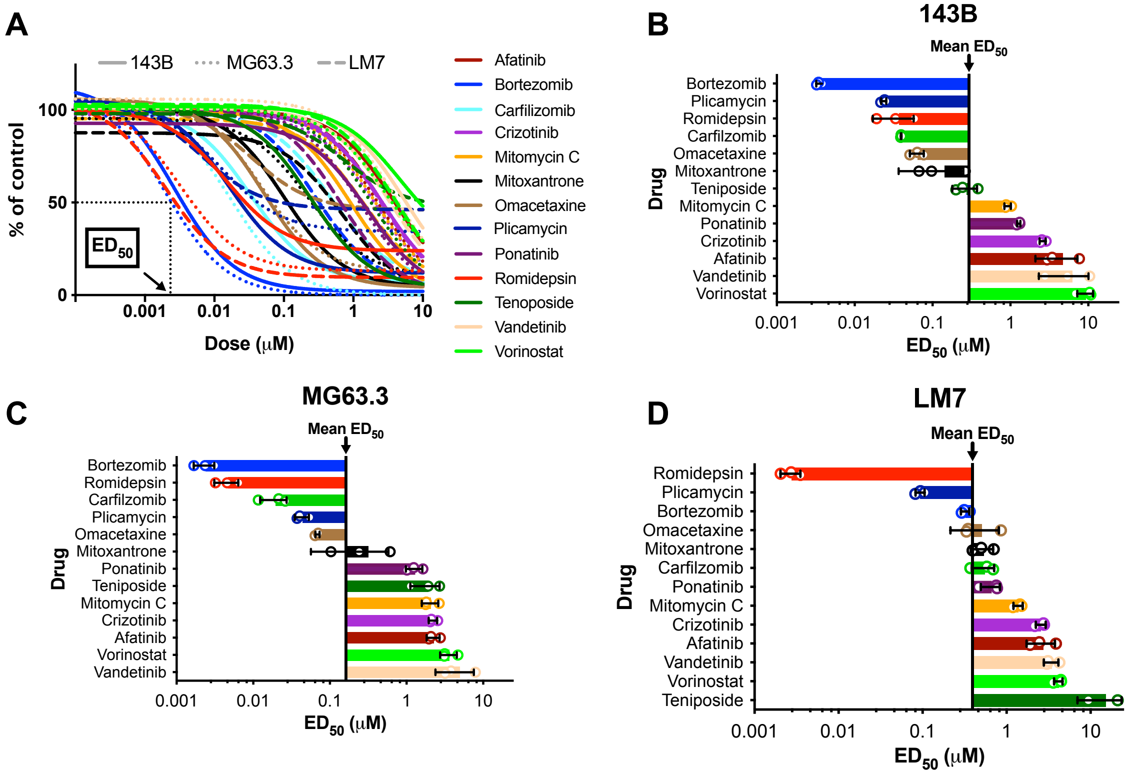


**Supplemental Figure S1. Dose Responses and ED50s from secondary screen.** (A) Dose responses (n=3 independent experiments) for 13 drugs included in secondary screen on sarcospheres derived from the highly metastatic cell lines. (B-D) Bar charts represent ED50s calculated from dose response curves shown in panel A. Overall mean ED50 is represented by vertical line. Error bars represent standard deviation.

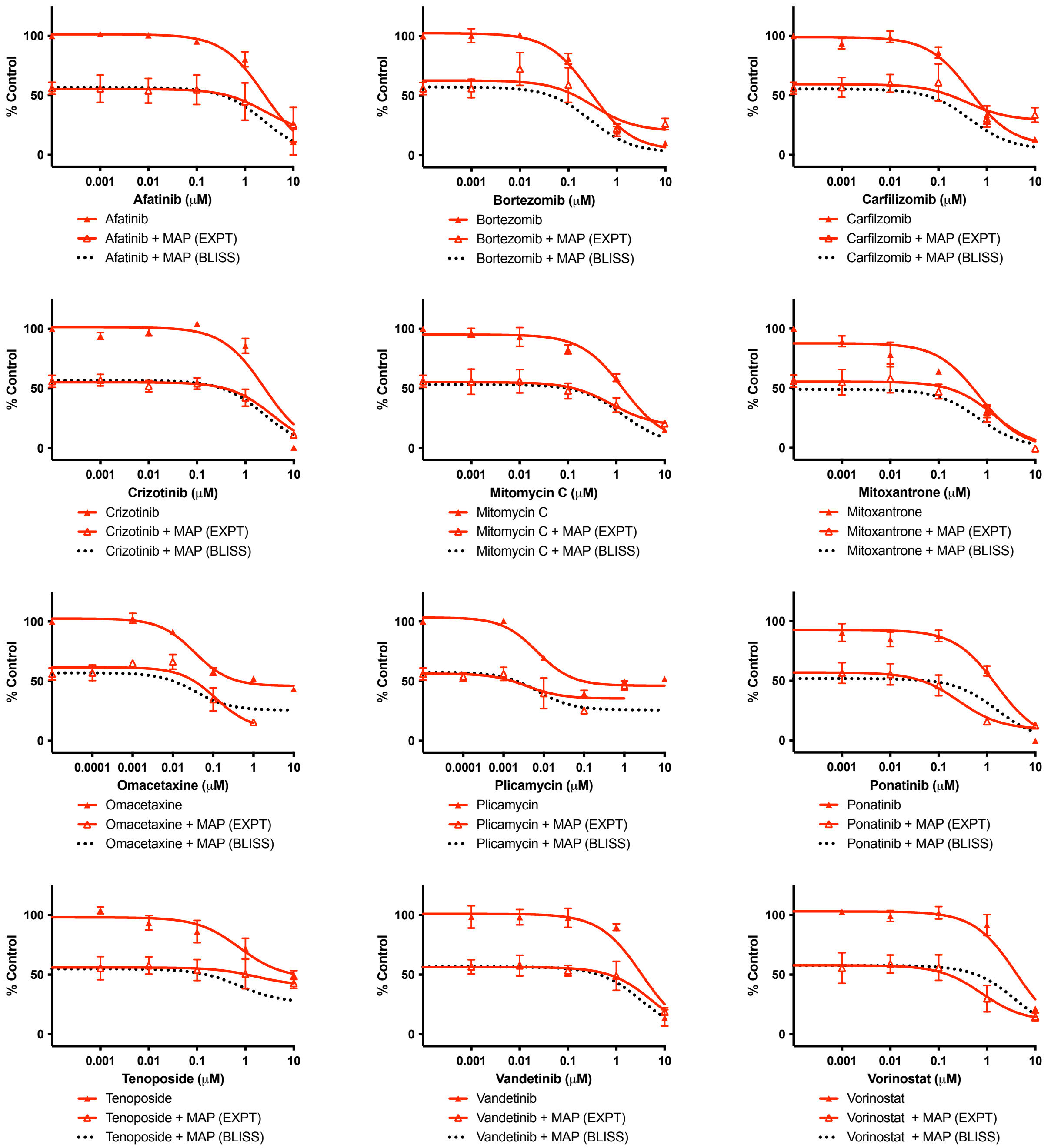


**Supplemental Figure S2. Dose Responses and Bliss Prediction of Additivity for LM7 sarcospheres in secondary screen.** Equivalent results for romidepsin are shown in Figure 3G**.** Data shows mean and standard deviation (n=3 independent experiments).

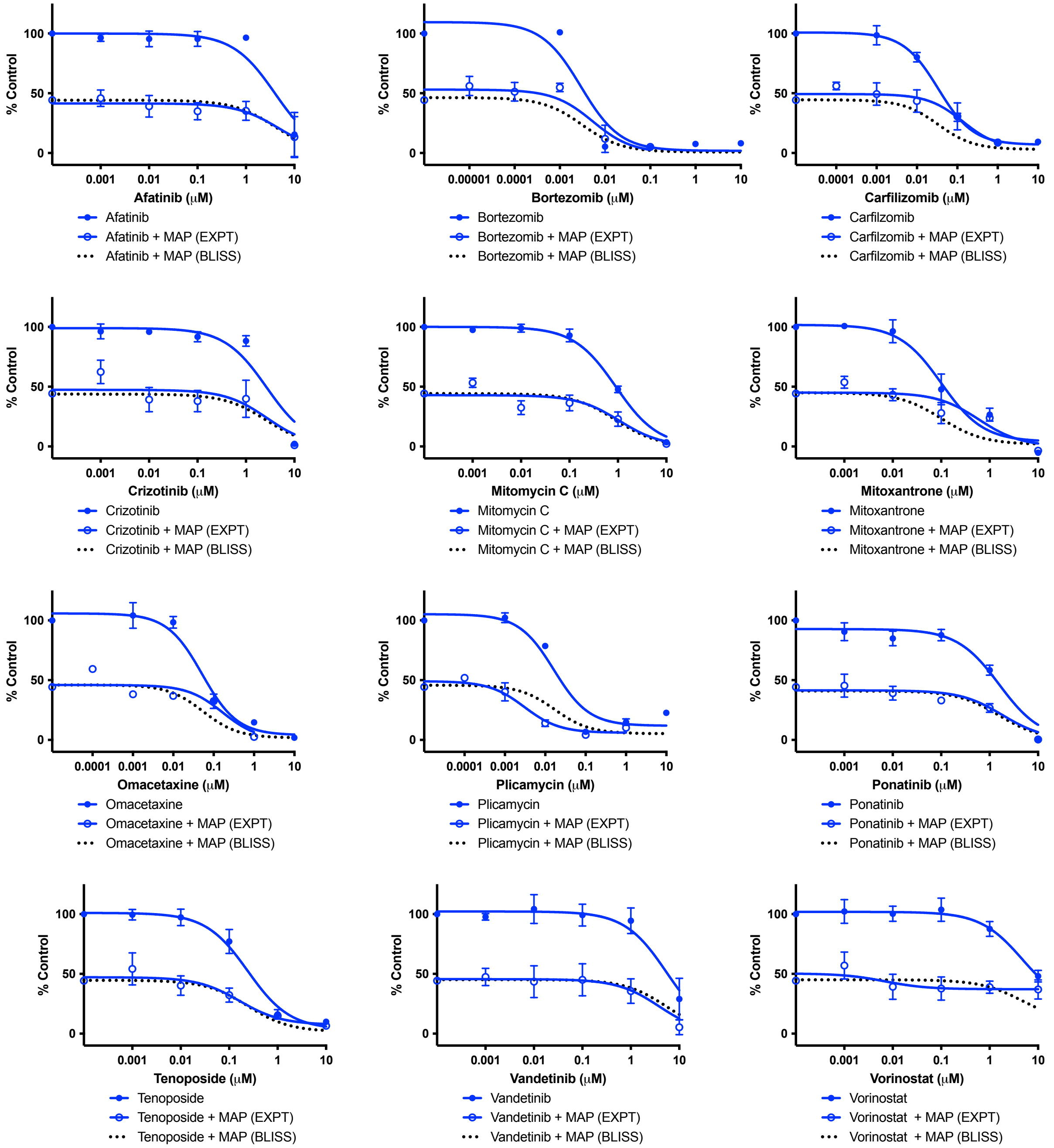


**Supplemental Figure S3. Dose Responses and Bliss Predictions of Additivity for 143B sarcospheres in secondary screen.** Equivalent results for romidepsin are shown in Figure 3E. Data shows mean and standard deviation (n=3 independent experiments).

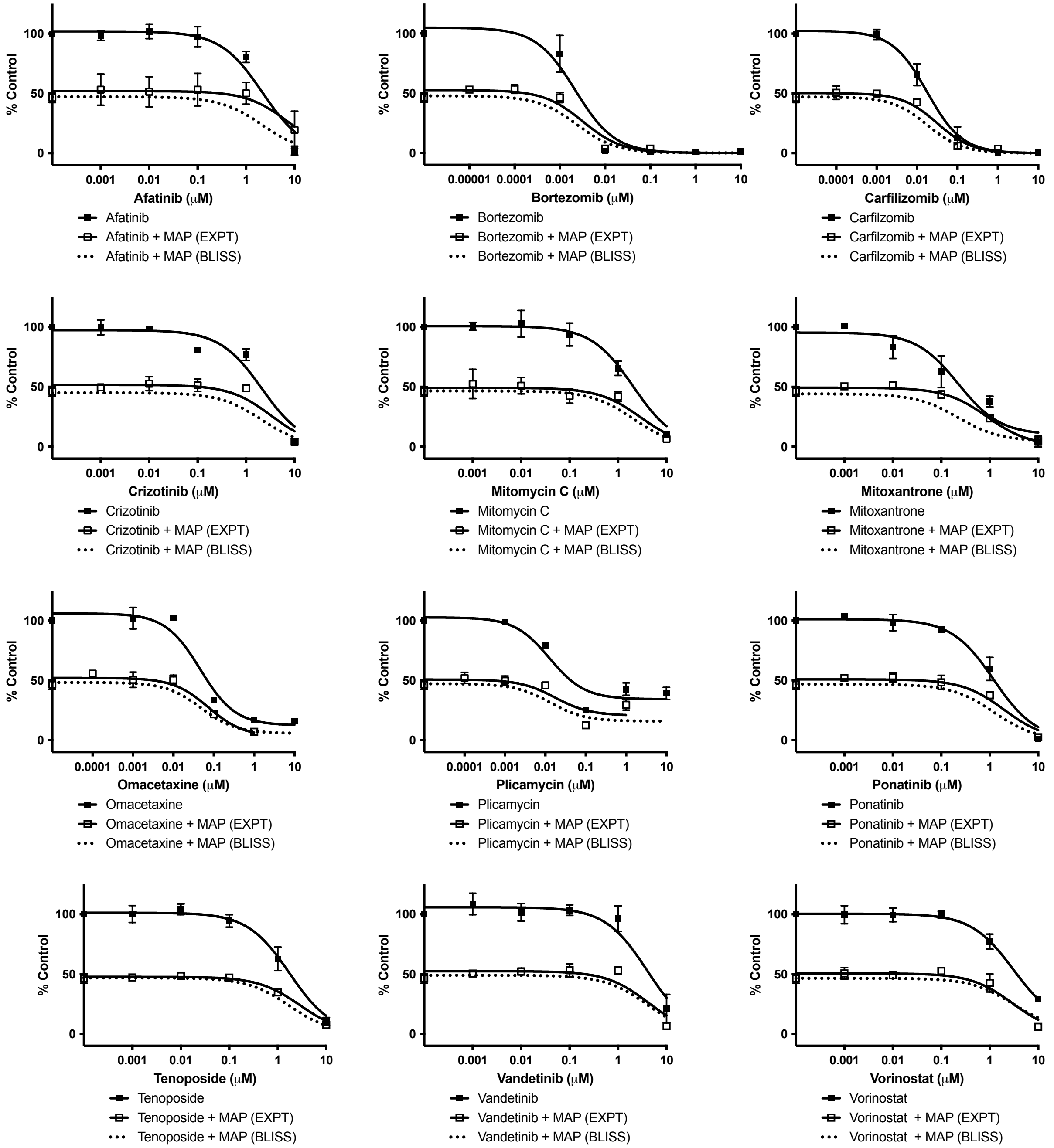


**Supplemental Figure S4. Dose Responses and Bliss Prediction of Additivity for MG63.3-GFP sarcospheres in secondary screen.** Equivalent results for romidepsin are shown in Figure 3F. Data shows mean and standard deviation (n=3 independent experiments).

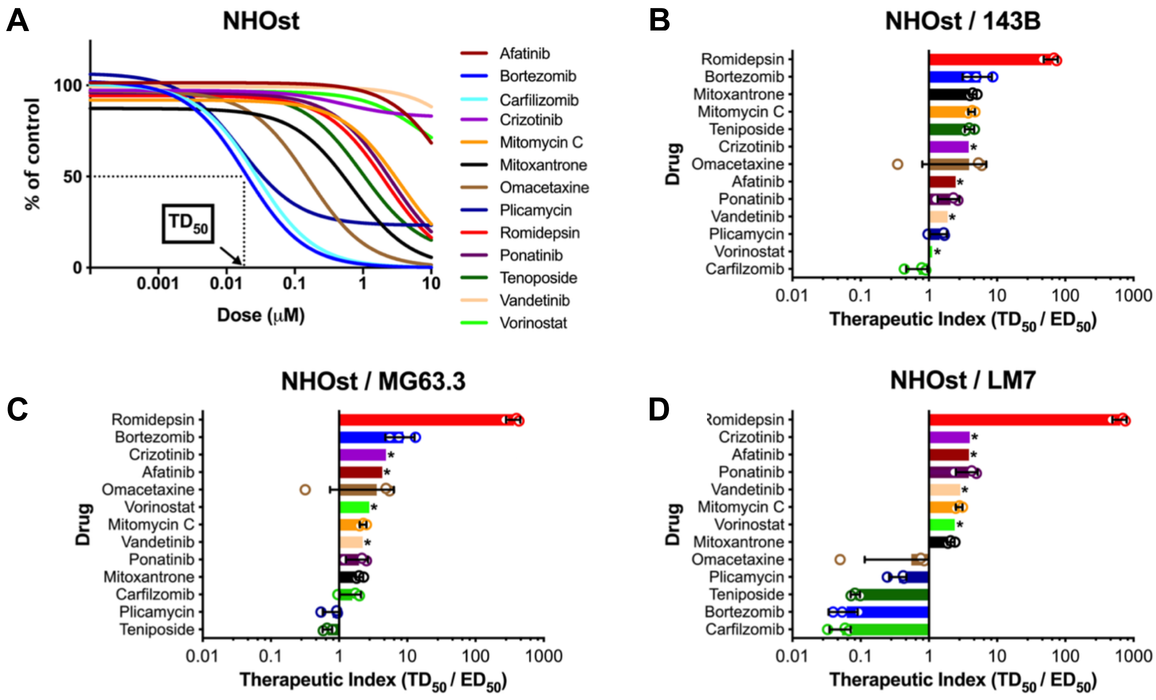


**Supplemental Figure S5. Toxicity and therapeutic indices on Normal Human Osteoblasts (NHOst).** (A) Dose responses for 13 drugs included in secondary screen on NHOst monolayers (n=3 independent experiments). (B-D) Bar charts represent therapeutic index (TD50/ED50 ratios) for 143B, MG63.3-GFP, and LM7 respectively for the 13 drugs. TD50s were determined from dose response curves in panel A and sarcosphere ED50s were determined from Supplementary Figure 1B-D. Error bars represent standard deviation. Asterisks indicate that 10uM, the highest dose that was tested, did not inhibit NHOst resazurin reduction by 50%.

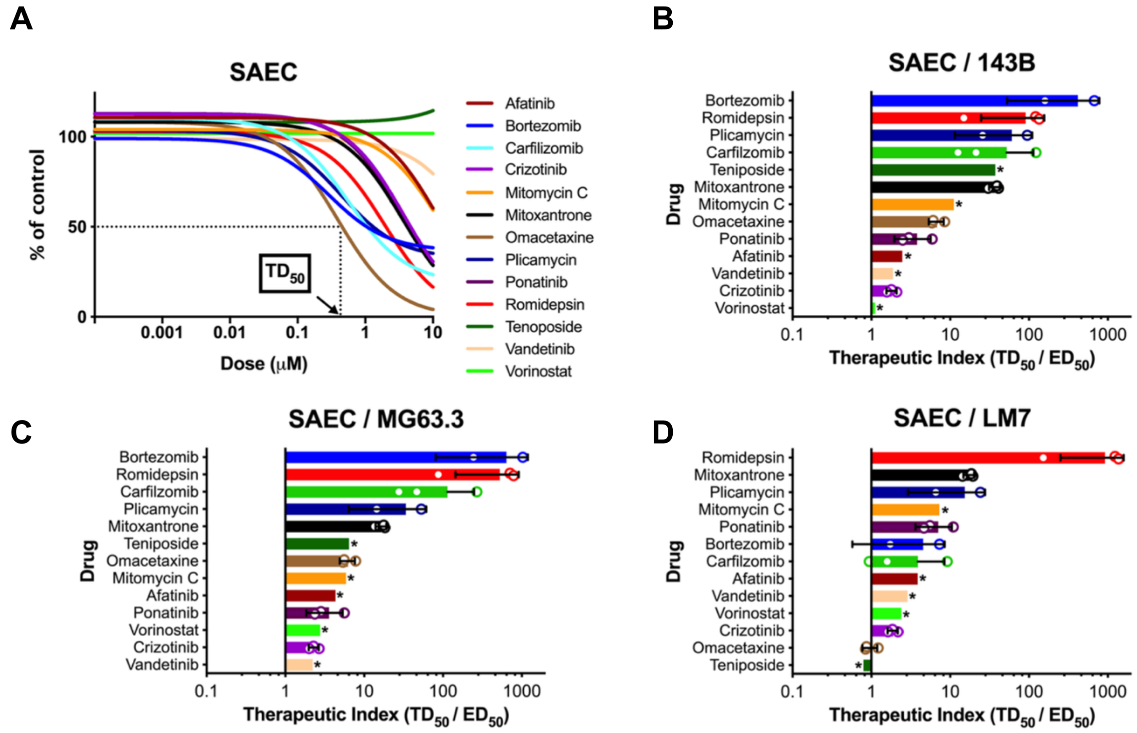


**Supplemental Figure S6. Toxicity and therapeutic indices on non-transformed human Small Airway Epithelial Cells (SAEC)).** (A) Dose responses for 13 drugs included in secondary screen on SAEC monolayers (n=3 experiments). (B-D) Bar charts represent therapeutic indices (TD50/ED50 ratios) for 143B, MG63.3-GFP, and LM7 respectively for the 13 drugs. TD50s were determined from dose response curves in panel A and sarcosphere ED50s were determined. from Supplementary Figure 1B-D. Error bars represent standard deviation. Asterisks indicate that 10uM, the highest dose that was tested, did not inhibit SAEC resazurin reduction by 50%.

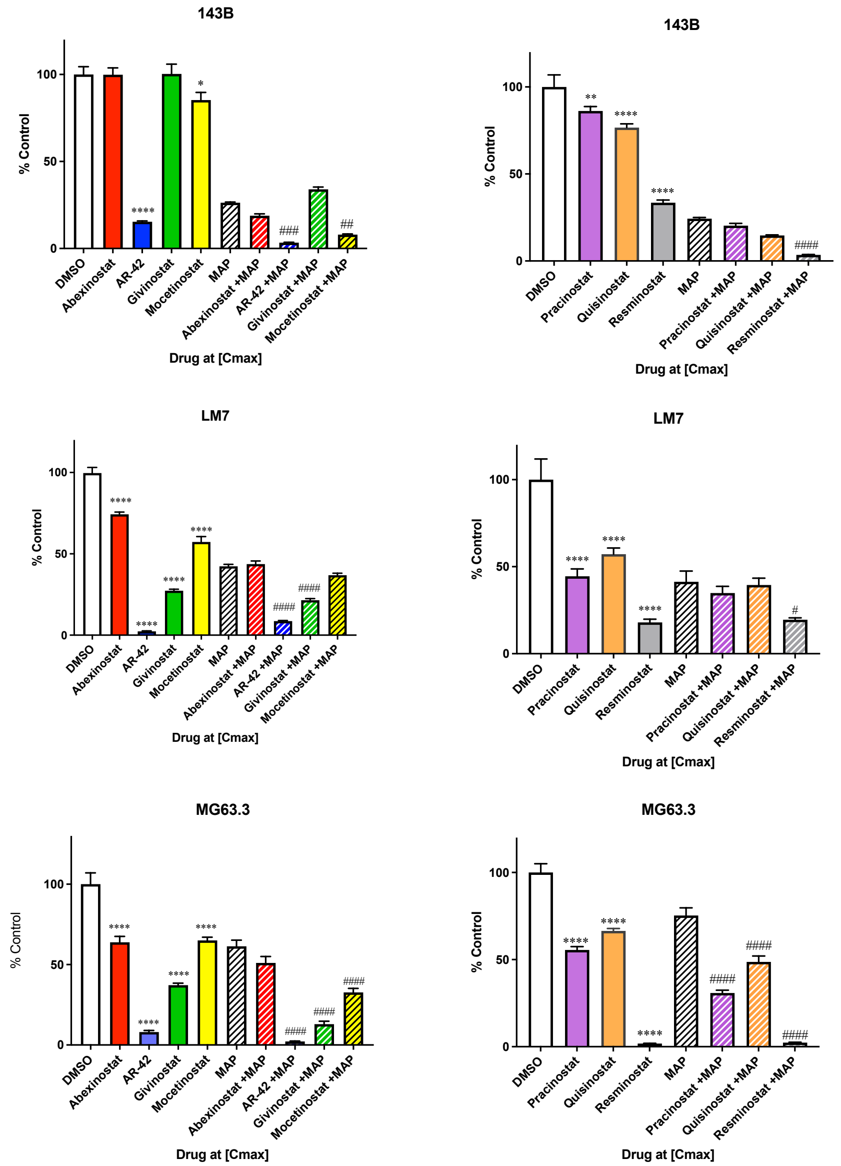


**Supplemental Figure S7 Screen of HDIs in clinical trials.** HDIs were tested at their Cmax with and without standard of care MAP to select HDIs for further study. HDIs were tested in two separate groups: abexinostat, AR-42, givinostat, and mocetinostat were in one group (A-C) and practinostat, quisinostat, and resminostat were in the other group (D-F). Sarcospheres were derived from the 143B (A-B), LM7 (C-D), and MG63.3-GFP (E-F) cell lines (n=3-5 independent experiments/cell line). Two-way ANOVA with Sidak's multiple comparisons test was used to compare each drug treated group with DMSO control (*p<0.05, **p<0.01, ***p<0.001, ****p<0.0001) or MAP control (#p<0.05, ##p<0.01, ###p<0.001, ####p<0.0001).

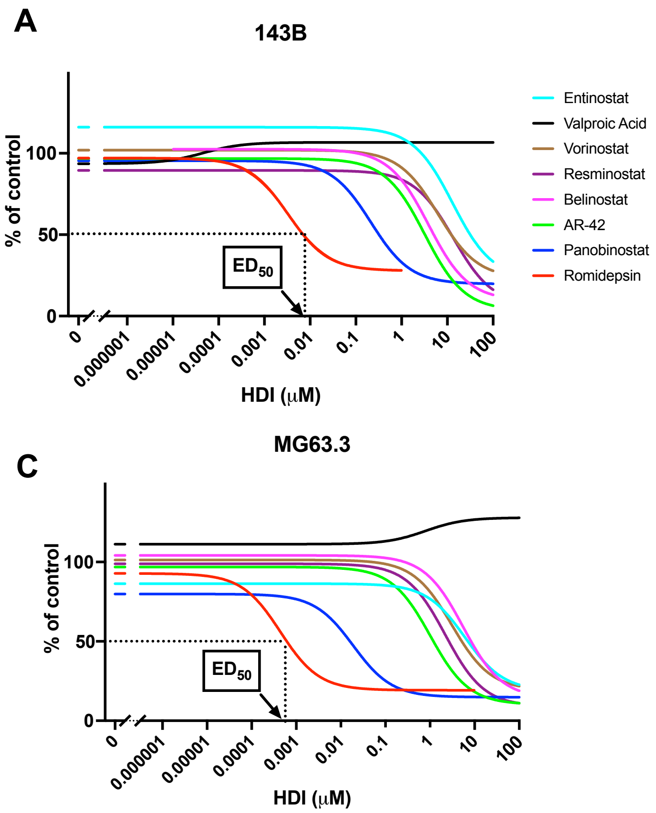


**Supplemental Figure S8. Dose Responses for HDIs on 143B, LM7, and MG63.3-GFP Sarcospheres.** (A-C) Dose responses (n=3 independent experiments) for HDIs on 143B, LM7, and MG63.3-GFP sarcospheres respectively.
